# Aberrant recursive splicing in a human disease locus

**DOI:** 10.1101/2025.08.14.666599

**Authors:** Philip M. Boone, Ricardo Harripaul, Rachita Yadav, Michael Grzybowski, Mahmoud K. Hanafy, Amanda C. Lee, Esther Y. Choi, Ryan L. Collins, Oksana Polesskaya, Nina Makhortova, Matthew O. Larson, Hakan Kayir, Yizhi Wang, Rodolfo A. Avila, Jude A. Frie, Amr Eed, Abdalla M. Albeely, Sunitha Vemuri, Samantha M Ayoub, John M. Lemanski, Daniel Ben-Isvy, Xuefang Zhao, Alba Sanchis-Juan, Maris Handley, Serkan Erdin, Celine de Esch, Kiana Mohajeri, Clementine Chen, Paulina Gonzalez Tovar, Monica Salani, Mariana Moyses-Oliveira, Derek J.C. Tai, Benjamin Currall, Christopher McGraw, Susan Slaughenhaupt, Ryan Doan, Dadi Gao, James F. Gusella, Sandra Sanchez-Roige, Jared W. Young, Jibran Khokhar, Aron M. Geurts, Abraham A. Palmer, Michael E. Talkowski

**Affiliations:** Center for Genomic Medicine, Massachusetts General Hospital, Boston, MA, US; Program in Medical and Population Genetics, The Broad Institute of MIT and Harvard, Boston, MA, US; Division of Genetics and Genomics, Boston Children’s Hospital, Boston, MA, US; Stanley Center for Psychiatric Research, The Broad Institute of MIT and Harvard, Boston, MA, US; Department of Neurology, Massachusetts General Hospital, Boston, MA, US; Department of Physiology, Medical College of Wisconsin, Milwaukee, WI, US; Department of Anatomy and Cell Biology, Western University, London, ON; Department of Psychiatry, University of California San Diego, La Jolla, CA, US; Human Neuron Core, Boston Children’s Hospital, Boston, MA, US; Division of Medical Sciences, Harvard Medical School, Boston, MA, US; Harvard Stem Cell Institute Flow Cytometry Core, Massachusetts General Hospital, Boston, MA, US; Department of Neurology, Boston Children’s Hospital, Boston, MA, US; Molecular Neurogenetics Unit, Massachusetts General Hospital, Boston, MA, US; Department of Genetics, Blavatnik Institute, Harvard Medical School, Boston, MA, US; Institute for Genomic Medicine, University of California San Diego, La Jolla, CA, US; Department of Medicine, Division of Genetic Medicine, Vanderbilt University, Nashville, TN, US

**Keywords:** Recursive splicing, *CADM2*, synaptic cell adhesion molecule 2, synCAM, ADHD, RNA-seq, induced neurons

## Abstract

Recursive splice sites are rare motifs postulated to facilitate splicing across massive introns and shape isoform diversity, especially for long, brain-expressed genes. The necessity of this unique mechanism remains unsubstantiated, as does the role of recursive splicing (RS) in human disease. From analyses of rare copy number variants (CNVs) from almost one million individuals, we previously identified large, heterozygous deletions eliminating an RS site (RS1) in the first intron of *CADM2* that conferred substantial risk for attention deficit hyperactivity disorder (ADHD) and other neurobehavioral traits. *CADM2* encodes a neuronally expressed cell adhesion molecule that has repeatedly been associated with ADHD and numerous similar traits. To explore the molecular impact of RS ablation in *CADM2*, we used CRISPR to model patient deletions and to target a smaller region (∼500 base pairs) containing RS1 in both human induced neurons (iNs) and rats. Transcriptome analyses in unedited iNs provided a catalog of *CADM2* transcripts, including novel transcripts that retained RS exons. Intriguingly, ablating RS1 altered the gradient of RNA abundance across the first intron of *CADM2*, decreased the level of *CADM2* expression, and impacted transcript usage. Decreased *CADM2* expression was reflected in reduced exon usage downstream of the RS1 site and global alteration to genes involved in neuronal processes including synapse and axon development. Given the scale of our analyses and the widespread association of *CADM2* with neurobehavioral traits, we sought to validate these findings using *in vivo* models and found that rodent models harboring *Cadm2* RS1 deletions exhibited significant changes in relevant behaviors and functional brain connectivity. In summary, our analyses demonstrate a functional role for RS as a noncoding regulatory mechanism in a gene associated with a spectrum of neuropsychiatric and behavioral traits.

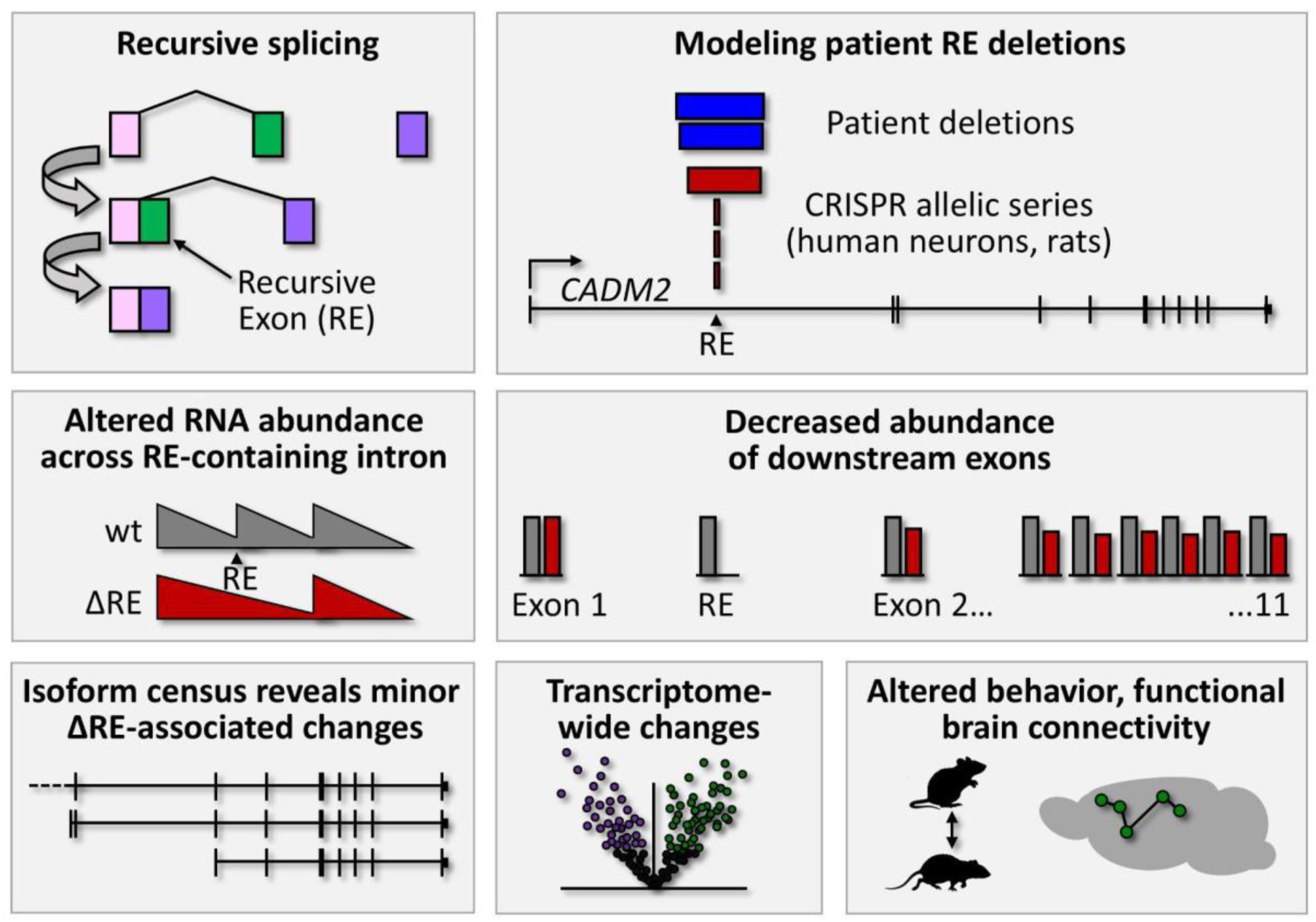
Graphical Abstract.

## Introduction

While mutations that alter canonical splicing are routinely surveyed in genetic association studies, one regulatory process yet to be implicated in human disease is recursive splicing (RS).^1–3^ The mechanism of RS, also called zero-exon splicing, was initially discovered in flies.^4–6^ It involves the removal of long introns from pre-mRNAs in two steps. An RS site (“ratchet site” in fly) divides the intron into two segments to be spliced out sequentially (Fig. 1f). The RS site initially functions as a 3′ (acceptor) splice site as the first intronic segment is removed. The RS site then contributes to a reconstituted 5′ (donor) splice site, facilitating the removal of the second intronic segment and the recursive exon (RE) from the maturing mRNA. RS has only recently been demonstrated in humans among a handful of long brain-expressed genes.^4,7^ These RS sites are evolutionarily conserved, and while initial studies in model organisms suggested that RS contributes to fidelity and diversity of splicing, this has yet to be confirmed in humans.^8–10^ Moreover, the role of RS in human health and disease is unknown.

**Figure 1.**
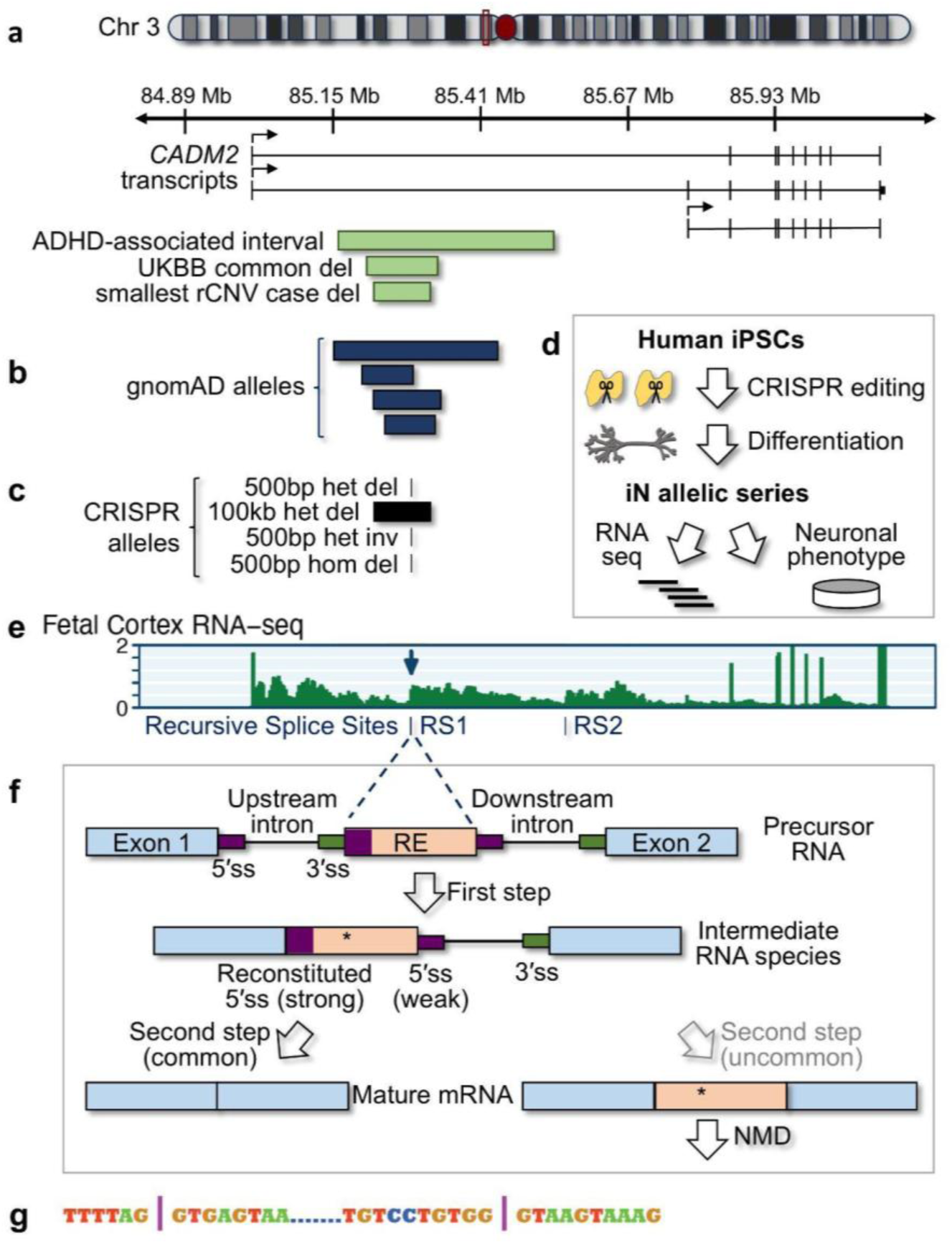
Identification and CRISPR modeling of an intronic *CADM2* recursive splice (RS) site deletion associated with hyperactivity and other neurobehavioral phenotypes. **a**. A case-control analysis of noncoding genomic rare copy-number variants (rCNVs), using array-based data, identified intronic deletions of *CADM2* recursive splice site 1 (RS1) in genome-wide significant association with hyperactivity (ADHD).^11^ ‘ADHD-associated interval’ is the 95% credible interval containing the ADHD rare deletion association from meta-analysis across sub-cohorts. ‘UKBB common del’ is the most frequent deletion allele among the largest rCNV data source, the UK Biobank. ‘Smallest case del’ is the smallest deletion in an individual with ADHD that encompasses *CADM2* RS1. See Table S3 for exact coordinates. GTEx transcripts and the local position on chromosome 3 (maroon box) are shown. **b**. In the present work, we identified rare overlapping intronic deletions in gnomAD, and All of Us (not shown), via WGS data. **c.** We generated an allelic series deleting or inverting RS1. See Table S6 for exact coordinates. Het, heterozygous. Hom, homozygous. Del, deletion. Inv, inversion. **d.** This allelic series was generated via CRISPR engineering in human induced pluripotent stem cells (iPSCs), which were then differentiated into glutamatergic neurons (iNs) for downstream functional genomic analyses. e. *CADM2* RS sites, which are marked in fetal cortex total RNA-seq read depth (from ^12^) by an uptick of abundance in the absence of an annotated exon. This “sawtooth” pattern of read depth is characteristic of recursive splicing. **f.** Recursive splicing in humans, modeled after (Cook-Andersen Nature 2015). RE, recursive exon. * Premature termination codon. ss, splice site. The RS site is the 3′ss at the left-hand side of the RE. NMD, nonsense-mediated decay. *CADM2* RS1, the first of two RS sites in *CADM2*, follows this characteristic architecture of human RS described in long, brain-expressed genes: It is present in the extremely long (767 kb) first intron of the gene, the site is followed by a recursive exon (RE) necessary for recognition of the RS site by splicing machinery but whose own ‘native’ 5′ splice site (5′ss) is weaker than the reconstituted 5′ss that is formed upon splicing of the exon 1 5′ss to the RS site,^13,14^ and the RE contains a premature termination codon (PTC) predicted to trigger nonsense-mediated decay if included in a mature transcript (^7^; Fig. S5). This architecture contrasts with the second RS site in *CADM2* (RS2) – not the focus of this study – which is overlapped by an alternative exon that can be included in mature transcripts when *CADM2* is driven by a minor promoter (^7,15^; Fig. S5). **g.** RE1 (between | marks) and flanking nucleotides.

We recently completed a study of genomic copy number variants (CNVs) in 950,278 individuals across 54 primarily neurological disease categories.^11^ A cluster of rare, large (∼100 kb), heterozygous deletions in the first intron of *CADM2*, which encodes a brain-expressed synaptic cell adhesion molecule, was collectively associated with risk of ADHD and other neurobehavioral traits. Intriguingly, common variants in and around *CADM2* have repeatedly been linked to ADHD and related phenotypes, including impulsivity and drug/alcohol use via genome-wide and phenome-wide association studies, highlighting *CADM2* as an important contributor to these phenotypes (https://www.ebi.ac.uk/gwas/genes/CADM2 and ^16–22^). Furthermore, *CADM2* is one of the few human genes known to utilize RS,^4,7^ and the intronic deletions from our CNV study remove an RS site from the gene without affecting its coding sequence. Here, we model alterations to RS in *CADM2*, demonstrating a functional role for RS and illuminating disease biology relevant to the molecular architecture of neurobehavioral traits.

## Results

### Further characterization of CADM2 recursive splice site deletions

In a recent survey of rare copy-number variants (rCNVs), we identified 163 genomic loci where rCNVs were significantly associated with one or more disease phenotypes.^11^ Twelve of these did not overlap any known protein-coding sequences. Among these 12 was a cluster of rare deletions localized to intron 1 of *CADM2* (chromosome 3p12.1; Table S3) and overlapping a previously identified *CADM2* RS site (RS1) (Fig. 1).^4,7^ These deletions were carried at a rate of ∼1:4,600 ADHD cases vs ∼1:20,000 control individuals, resulting in a strong association with ADHD (meta-analysis natural logarithm odds ratio (lnOR) 3.44; 95% CI 3.10-3.76; p=2.2e-5) and nominal association with bipolar disorder (p=4.31e-3, lnOR 1.48) and other related phenotypes (Table S8).

The alleles identified in our discovery cohort were limited by methodological considerations of microarray-based rCNV analyses (e.g., size >100 kb).^11^ Thus, to ascertain a population allelic spectrum of intronic deletions encompassing *CADM2* RS1, we genotyped structural variants (SVs) from large cohorts sequenced by whole genome sequencing (WGS). WGS data from 63,046 individuals in gnomAD^23^ revealed four distinct intronic deletion alleles of the *CADM2* RS1 site (Table S9; Figs. 1b, S14). These deletions are similar in size and collective allele frequency (AF 7.14e-5, or ∼1/14,000 alleles) to the variants in our discovery cohort. Each gnomAD deletion is specific to a genetic ancestry group. None of these SVs is tagged by a common (AF ≥ 1%) SNP in gnomAD (r^2^ < 0.8), but two are tagged by one or more rare SNPs also specific to that ancestry group and of equivalent allele frequency to the respective deletion (Table S9). We also investigated SV calls from WGS of 84,247 individuals in All of Us, representing an ancestrally more diverse dataset than gnomAD.^24^ This identified five deletion alleles removing RS1, two of which matched gnomAD alleles and three of which were novel (Table S9). No *CADM2* RS1 deletions were found among GTEx SV calls from 600 subjects.^25^ We also assessed gnomAD for single-nucleotide variants (SNVs) affecting the *CADM2* recursive exon 1 (RE1) and flanking sequence. Several rare SNVs were present in the exon but none affected the conserved 5′ splice site (ss) or 3′ss (Table S9, Fig. S14).

### In vitro models of CADM2 intronic deletion show disrupted recursive splicing

We sought to extensively characterize the molecular and functional consequences of ablating RS in *CADM2*. We modeled intronic *CADM2* deletions in human induced pluripotent stem cells (iPSCs; Table S1, Fig. 1c-d) and differentiated these into neurogenin 2 (*Ngn2*) induced glutamatergic neurons (iNs) to identify how the deletions impact recursive splicing, *CADM2* expression, the broader transcriptome, and neuronal function. Specifically, we used CRISPR to replicate patient deletions from our association analysis by engineering heterozygous 102 kb deletions (denoted “het 100 kb del”) and targeted heterozygous or homozygous 599 bp deletions (“het 500 bp del” and “hom 500 bp del”, respectively) or inversions (“het 500 bp inv”), containing the RS1 site and RE1. *CADM2* was robustly expressed in differentiated iNs as measured by RNA sequencing (RNA-seq; average TPM 49.79) and more modestly expressed in iPSCs (average TPM 14.56) (Fig. S7a).

In both iPSCs and iNs, deletion or inversion of RS1 ablated recursive splicing in a zygosity-dependent manner (Figs. 2, S4, S9, S8g). As total RNA-seq reads spanning the exon 1 to RE1 junction are rare (Fig. S13a), we quantified this junction via qPCR; splicing of exon 1 to RE1 was eliminated in hom 500 bp del iNs and decreased by 44-47% in heterozygous iNs (all p=<1e-6 by Tukey post-hoc HSD test) (Fig. 2a), confirming our mutant human cell lines as robust models of RS disruption in the pluripotent stem cell and postmitotic neuronal states.

**Figure 2.**
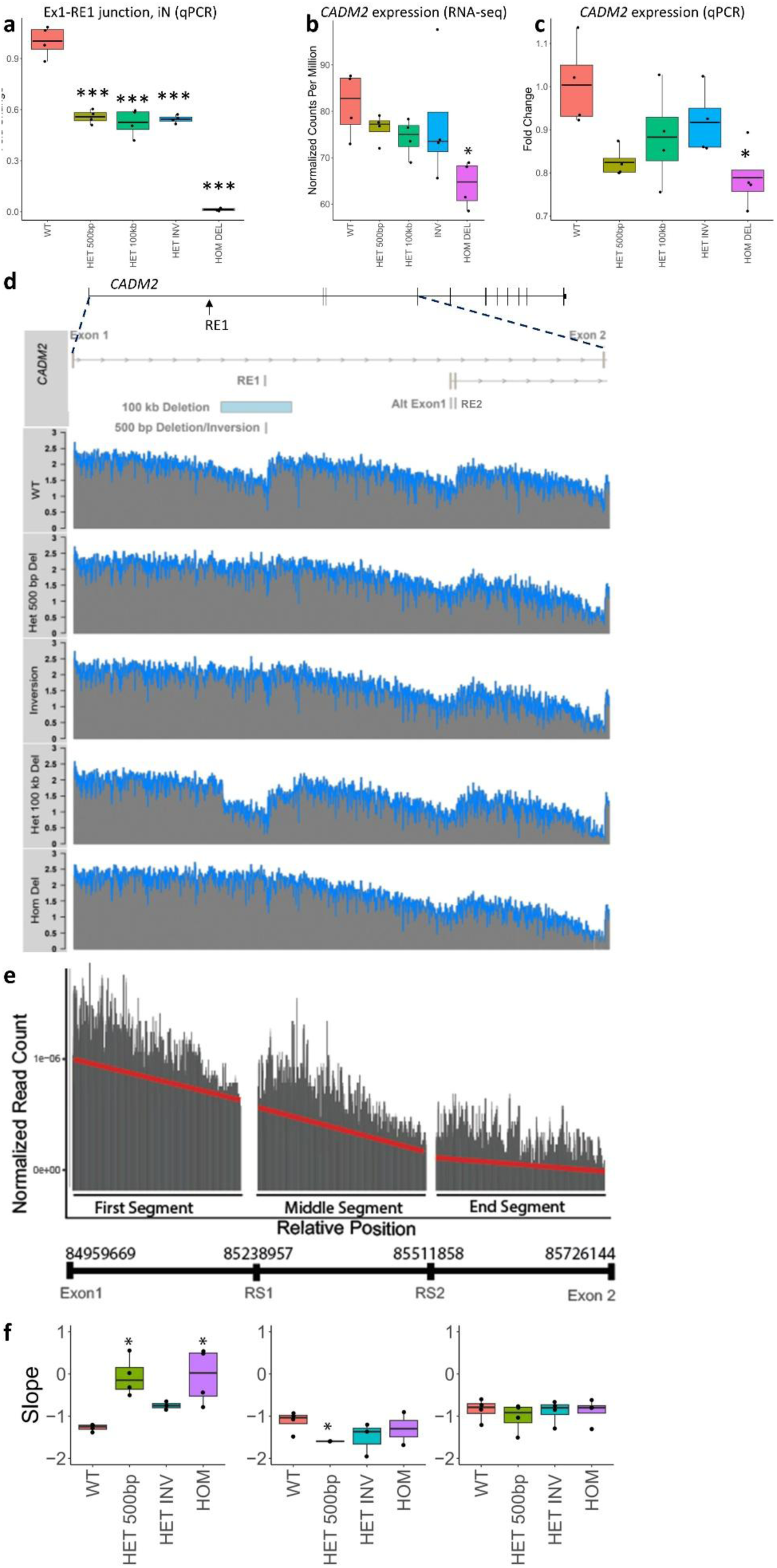
CRISPR *CADM2* RS1 site deletion in human induced neurons (iNs) ablates recursive splicing, alters the pattern of intronic mRNA abundance, and decreases *CADM2* expression. **a**. qPCR using primers amplifying from exon 1 to RE1 confirms that RS1 deletion or inversion ablates recursive splicing in a zygosity-dependent manner, with a decrease of 44-47% in het and of 99% in hom lines, compared to wt (one-way ANOVA p=1.53e-14; all comparisons to wt p<1e-6 by Tukey’s HSD test). Normalization gene is *GUSB*. **b.** Total RNA-seq demonstrates that RS1 site deletion or inversion decreases the overall expression of *CADM2*, in a zygosity-dependent manner, by 23% in hom 500 bp del iNs (FDR=7.4e-3, Wald’s test) as compared to wt. **c**. qPCR using primers amplifying between constitutive exons 6 and 8 confirms that RS1 site deletion decreases expression of *CADM2* in a zygosity-dependent manner, by 8-18% of wt in het lines and by 21% of wt in hom 500 bp del (one-way ANOVA p=0.0021 and individual comparisons to wt significant only for hom 500 bp del iNs with p=0.018 by Tukey’s HSD test; with hets combined, one-way ANOVA p=0.0082 and comparisons to wt significant for hom 500 bp del (p=0.0066) and combined hets (p=0.046) by Tukey’s HSD test). Normalization gene is *GUSB*. **d.** Normalized total RNA-seq coverage (log_2_ reads/bp) mapped across *CADM2* intron 1 shows the sawtooth pattern of read depth corresponding to known waypoints of recursive splicing in this gene (RS1, RS2). RS1 deletion or inversion alters the sawtooth pattern. Blue represents smoothened data. See also Fig. S16. **e.** Segmentation of the sawtooth pattern over *CADM2*’s first intron for one wt sample. The first segment is from exon 1 to RS1, the middle segment is from RS1 to RS2, and the end segment is from RS2 to exon 2. The red line represents the read count slope within a segment. Chr3 coordinates of the segment boundaries are shown. **f.** Read count slopes (normalized read depth/cumulative relative position), by genotype, within the segments defined in (e). Ablation of RS1 flattens the slope in segment 1, has no effect in segment 2 except for in het 500 bp del (see text), and has no effect on the slope in segment 3 (First segment: hom 500 bp del: p=3.4e-2, het 500 bp del p=1.2e-2; Middle segment: het 500 bp del p=9.7E–3; all by Student’s t-test). Het 100 kb del was removed from the analysis on account of removing a large part of the intron. Het, heterozygous. Hom, homozygous. Del, deletion. Inv, inversion. Significance markings are * p<0.05, ** p<0.005, *** p<0.0005. (a) and (d) are standard boxplots. See Fig. S9 for similar data from iPSCs.

### Ablation of CADM2 RS1 alters the gradient of RNA abundance across the intron

Recursive splicing is expected to generate sequential pre-mRNA intermediates as the splicing machinery removes segments of the intron. The collection of nascent transcripts results in a gradient of higher intronic read coverage at each segment’s 5’ end than its 3’ end, owing to co-transcriptional splicing occurring relatively quickly when the transcript reaches the next segment.^26^ Prior studies have demonstrated this pattern of total RNA-seq read depth over introns containing RS sites, with higher depth immediately following exon 1, the RE(s), and exon 2, and tapering off to lower depth as the RE(s) and exon 2 are approached, creating a characteristic “sawtooth” pattern.^4,7^ We observed this intronic pattern in *CADM2* in unedited (denoted “wild type” or “wt” below) iPSCs and iNs (Fig. 2b, Fig. S9d).

Deleting RS1 visibly disrupts the recursive sawtooth pattern (Fig. 2b); specifically, the read depth abundance does not decay as the RS1 site is approached. To quantify changes to this pattern in RS1-edited lines, we calculated the slopes of total RNA-seq read coverage along the first intron within the three segments separated by REs (i.e., points of the saw): exon 1 to RE1; RE1 to RE2; and RE2 to exon 2 (Fig. 2e-f). The wt iN samples had slopes of –1.3 (first segment), –0.65 (middle segment), and –0.18 (end segment). Deleting RS1 led to changes in the slopes of the first and middle segments, but not the end segment (Fig. 2f). Compared to wt iNs, the first segment showed a flattening of the slope toward zero in the hom 500 bp del (slope=-0.05; p=6.0e-3; two-sided t-test) and het 500 bp del (slope=-0.06, p=5.2e-3) models. Of note, the het 100 kb del removes a large part of the intron, obscuring potential changes to read depth slope and precluding a slope comparison to wt. Analysis of the middle segment showed a mild slope steepening for het 500 bp del (slope=-1.85, p=9.72e-3), apparently as the result of a higher starting read depth at the beginning of the segment (Fig. S16). The third segment was unchanged in slope among the genotypes. iPSCs demonstrate similar visible changes to the sawtooth pattern of total RNA-seq read depth over *CADM2* intron 1 (Fig. S9d); when quantified this showed a similar pattern of flattening of the first segment’s slope (significant for hom 500 bp del, p=3.27e-2) (Fig. S9e).

### Deletion of CADM2 RS1 decreases overall CADM2 expression

In a first assessment of potential disease mechanisms, we evaluated the effect of RS1 deletion on overall *CADM2* expression (Figs. 2b-c, S9b-c). Total RNA-seq of iNs showed a 23% decrease of *CADM2* expression due to homozygous RS1 deletion (false discovery rate (FDR) =7.4e-3 by Wald’s test) and suggested intermediate decreases (8-12%; not individually significant) in all heterozygous genotypes (Fig. 2b). This result was confirmed by qPCR of constitutive *CADM2* exons, showing decreased expression (21%) in hom 500 bp del iNs (p=0.019 by Tukey post-hoc HSD test) and again suggesting mild decreases in all heterozygous genotypes (8-18%; not individually significant, but significant when all hets are combined (p=0.046)) (Fig. 2c). Junction reads across a constitutive exon junction (ex7-8) additionally confirmed a decrease in mutant lines (Fig. S13c).

Total RNA-seq of iPSCs, which exhibited variability potentially attributable to lower expression at this locus, showed a 62.5% decrease of overall *CADM2* expression in hom 500 bp del iPSCs (FDR=5.9e-5 by Wald’s test) (Fig. S9b). By qPCR, *CADM2* expression decreased by 59% in hom 500 bp del iPSCs (Tukey post-hoc HSD test p<1e-6), and by 5-28% in heterozygous iPSCs (significant for 500 bp del (p=8e-3) and 500 bp inv (p=8e-5), as well as for when all hets are combined (p=1.1e-2)) (Fig. S9c).

### Deletion of CADM2 RS1 decreases the expression of downstream exons

We next assessed *CADM2* exon usage among total RNA-seq data and compared among genotypes (Fig. 3e; Table S12). As expected, edited iNs showed decreased usage of RE1; its inclusion was completely absent in the hom 500 bp del (FDR=4.38e-2; t-test) and decreased by an average of 51% in the heterozygous models (p=2.24e–2; t-test) compared to wt. We also identified decreased expression of all highly expressed exons downstream of RE1, in hom 500 bp del iNs: exon 3 (21% reduction), 4 (24%), 5 (21%), 6 (17%), 7 (26%), 8 (29%), 10 (31%) and 11 (31%). The average decrease across these exons was 25%, closely mirroring changes to overall *CADM2* expression and suggesting a causal relationship between the two. Although not individually significant, heterozygous iNs also demonstrated decreased expression of exons downstream of RS1 (average 9% decrease). iPSCs also exhibited decreased expression of RE1 and exons downstream of RE1 in mutant genotypes (Fig. S8f). *CADM2*’s alternative exons (alt exon 1, exon 2, and exon 9), which if included versus excluded are predicted to encode slightly different proteins (Fig. S8b-c), were not differentially expressed in edited iNs; however, their low expression may limit statistical power to detect a change (Fig. 3e). *CADM2*’s annotated antisense transcripts (*CADM2-AS1* and *CADM2-AS2*) were also not differentially expressed (Fig. S8d-e).

**Figure 3.**
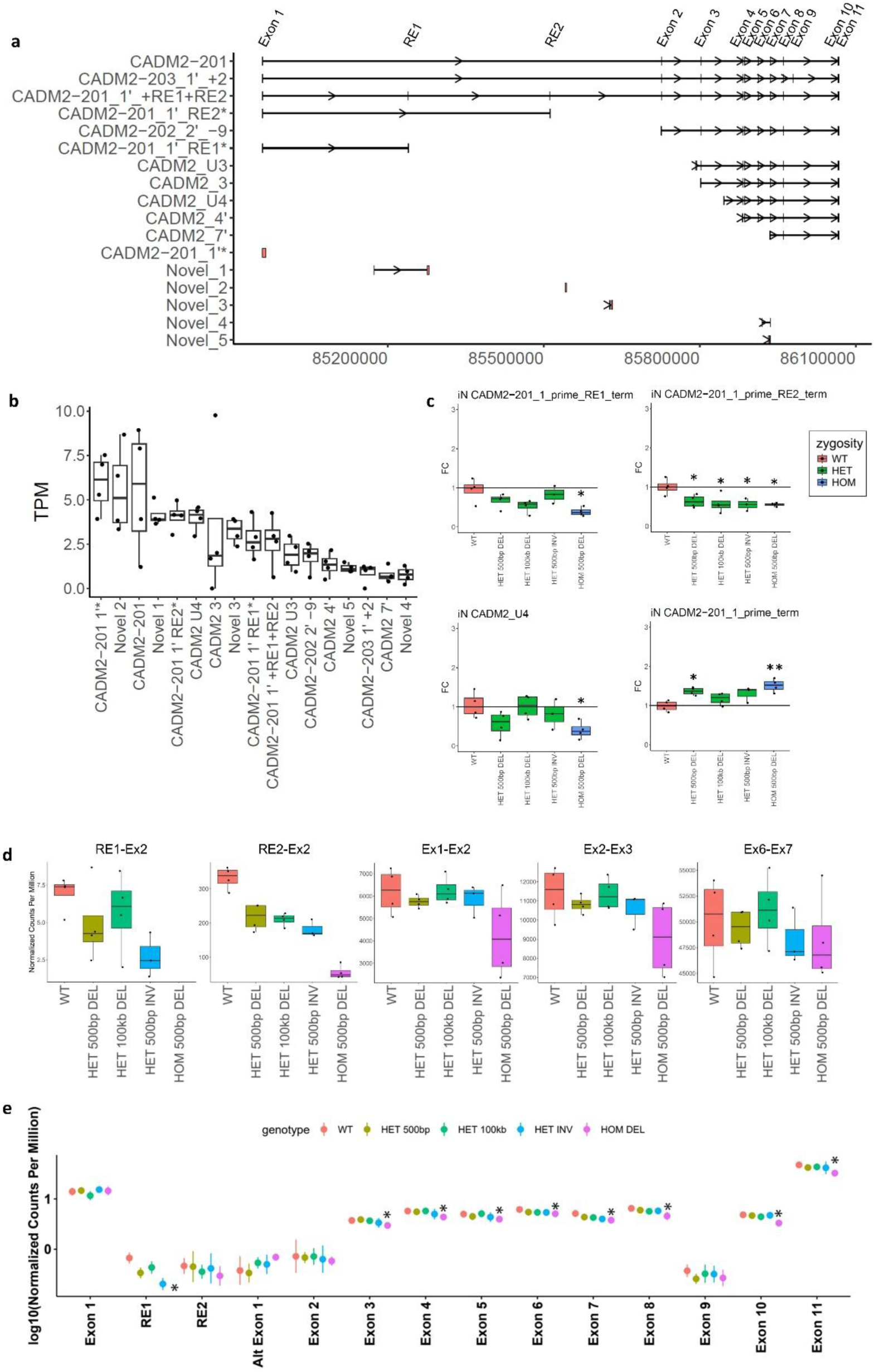
Ablation of *CADM2* RS in human iNs alters *CADM2* transcript abundance, splice junctions, and exon expression. **a.** *CADM2* transcripts identified by *de novo* transcript reconstruction combining total RNA-Seq and Cap-Seq data in iNs. Similarity to Ensembl transcripts is indicated by names (e.g. CADM2-201), with differences denoted as:’, differs from Ensembl exon definition; *, truncated at this exon; +, additional exon; –, missing exon; U, starts upstream of stated exon. Complete lack of overlap with Ensembl transcripts is denoted “Novel.” **b.** Normalized counts of reconstructed transcripts in wt iNs. See Table S13 for data. **c.** Standard boxplots showing fold change of selected transcripts’ TPM in iNs (see Fig. S19 for boxplots of all transcripts). This indicates that there are isolated effects on transcript choice from ablation of RS in *CADM2*. Statistics are two-sided t-tests between the wt and edited cell line fold changes. * p<0.05, ** p<0.005, *** p<0.0005. **d.** Genotype-dependent alterations in the abundance of junctions involving REs (RE1-Ex2, RE2-Ex2), without substantial changes in splicing between either early (Ex1-Ex2, Ex2-Ex3) or later (Ex6-Ex7) coding exons. Data are iN Cap-Seq junction reads, normalized to total reads (which are highly enriched for the *CADM2* locus). **e.** *CADM2* exon abundance via *DEXSeq*, using total RNA-Seq data. Data are read counts from iNs, normalized to library size. Exon 1 follows the definition of “exon1_long_full” in Table S4. The low abundance of RE1 and RE2, alt exon 1, exon 2, and exon 9 are visible. RS1 deletion results in mild decreases of exon abundance of exons downstream of RS1 that are of similar magnitude to, and potentially explain, the decrease in overall expression of *CADM2*. See text and Table S12 for statistics; displayed here is FDR. Figure S8 displays a similar pattern in iPSCs. * FDR<0.1.

### De novo CADM2 transcript reconstruction identifies novel and annotated transcripts

In addition to analyses of exon representation, we investigated the abundance of specific *CADM2* transcripts among edited iNs. GTEx/Ensembl transcripts are based on sequencing mature RNA (mRNA), making it challenging to quantify intermediate RNA molecules (i.e. before splicing is complete) including those generated via the process of recursive splicing (https://gtexportal.org/home/). To comprehensively catalog *CADM2* transcript diversity and explore transcript-specific changes associated with rCNVs altering RS1, we first augmented our total RNA-seq data by generating targeted RNA-seq data (Cap-Seq) at substantially higher depth using baits designed against known features of *CADM2*. We then performed a *de novo* transcript reconstruction (Methods) at the *CADM2* locus using both total RNA-seq and Cap-Seq data from our iN models (n=20). Briefly, this approach identifies reads within the *CADM2* region to generate representations of completely or partially spliced transcripts and maps them onto the exonic structure of *CADM2* (Table S4). Other parameters of note were that 80% of exons passed a counts per million filter (FPKM>10e-6) and that transcripts must occur in >50% of samples per genotype. This identified 17 transcripts (Fig. 3a). Most exhibited similarity with previously documented isoforms (CADM2-201, CADM2-202) – though in many cases with novel features – while five involved completely novel splice junctions and exons.

A unique aspect of these assembled transcripts is that one full-length transcript (“CADM2-201_1’_+RE1+RE2”) and two partial transcripts (“CADM2-201_1’_RE2*” and “CADM2-201_1’_RE1*) included one or both of the gene’s REs (RE1, 73 bp; RE2, 42 bp). The presence of these transcripts is supported by specific junction reads (e.g. Figs. S4b and S13b). The inclusion of REs in the full-length reconstructed transcript CADM2-201_1’_+RE1+RE2 could reflect a pre-mRNA, from which the REs will later be removed, or a mature but non-productive transcript that will be degraded by nonsense-mediated decay on account of the REs containing stop codons (Figs. S5, S6). Furthermore, its presence may indicate the ability of these RE’s 5’ splice sites to participate in splicing.

### Deletion of CADM2 RS1 affects expression of specific CADM2 transcripts

We quantified the *de novo* reconstructed *CADM2* transcripts in iNs (Fig. 3b), then compared the expression of these transcripts in RE1-edited and wt lines (Fig. 3c, Fig. S19). We identified genotype-dependent expression changes for seven of 17 transcripts. As expected, the transcript terminating in RE1 (CADM2-201_1’_RE1*) was significantly decreased in hom 500 bp del iNs (p=2.7e-2). We also identified decreased expression of the transcript terminating in RE2 (CADM2-201_1’_RE2*) in hom 500 bp del (p=1.8e-2), het 500 bp del (p=3.1e-2), het 500 bp inv (p=2.2e-2), and het 100 kb del (p=3.6e-2). Furthermore, a C-terminal transcript starting upstream of exon 4 (CADM2_U4) was decreased in expression in hom 500 bp del (p=2.1e-2). Interestingly, this result suggests downstream effects hundreds of kb from the deleted intronic RS1, affecting transcript usage. Finally, a transcript involving intron retention after exon 1 (CADM2-201_1’*) was increased in expression in hom 500 bp del (p=3.4e-3) and het 500 bp del (p=6.0e-3). Summing all reconstructed transcripts together allowed us to further confirm an overall decrease in *CADM2* expression upon deleting RE1 in iNs (Fig. S8h).

We also capitalized on the power of Cap-Seq to detect rare splice junctions in the *CADM2* locus in iNs. This demonstrated genotype-dependent alterations in the abundance of junctions (normalized to library size) involving REs (RE1-Ex2, RE2-Ex2), without substantial changes in splicing between either early (Ex1-Ex2, Ex2-Ex3) and later (Ex6-Ex7) coding exons (Fig. 3d).

### Deletion of CADM2 RS1 causes global transcriptomic effects centering on neuronal pathways

To identify whether *CADM2* RS1 deletion causes global transcriptomic effects, we performed differential gene expression analysis on each edited iN genotype compared to wt lines (Fig. 4; Table S10). The hom 500 bp del yielded 1,639 differentially expressed genes (DEGs) at p < 0.05. The het 100 kb del displayed 2,161 DEGs, the het 500 bp inv displayed 442 DEGs, and the het 500 bp del model displayed 259 DEGs. Comparisons of DEG sets among the edited iN genotypes demonstrated significant overlap (Fig. 4a), suggesting these DEGs are robust downstream effects of *CADM2* RS ablation. For example, comparison between the het 100 kb del and hom 500 bp del models showed significant sharing of DEGs (n=334, hypergeometric overlapping p=3.23e-26). At a higher DEG significance cut-off (FDR < 0.1), the het 100 kb del (327 DEGs) and hom 500 bp del (495 DEGs) iNs still showed significant DEG sharing (n=38; hypergeometric overlap p=2.97e-13).

**Figure 4.**
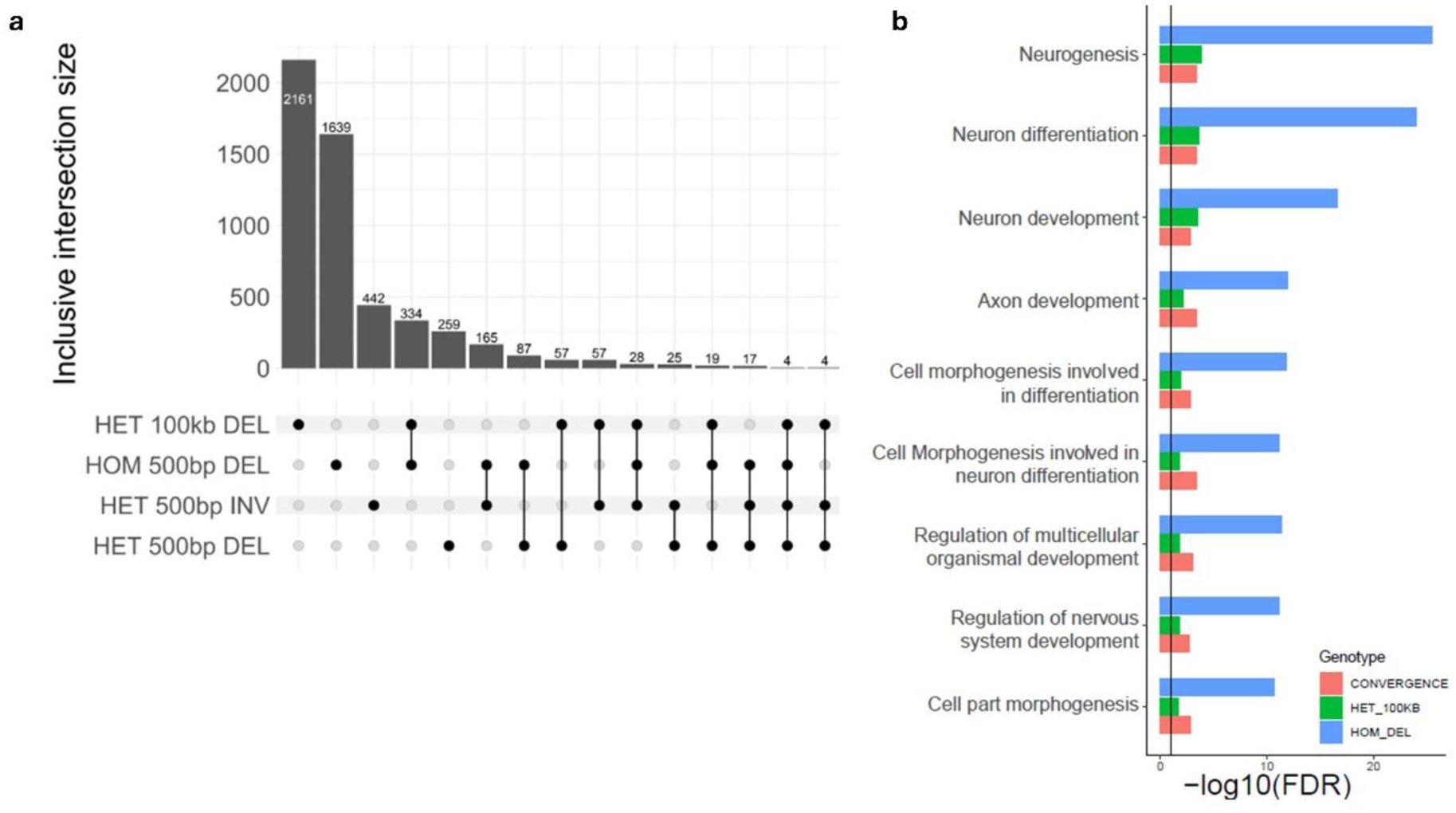
Deletion of *CADM2* RS1 in iNs results in transcriptomic disturbances across many genes representing specific neuronal/developmental pathways. **a**. Differentially expressed genes (DEGs) by genotype (comparisons to wt). Most two-way overlaps among DEG sets are significantly enriched (hom 500 bp del and het 100 kb del hypergeometric p=3.23e-26; hom 500 bp del and het 500 bp del p=4.26e-29; hom 500 bp del and het 500 bp inv p=8.49e-56; het 500 bp del and het 100 kb del p=1.88e-5; het 500 bp del and het 500 bp inv p=3.21e-8), attesting to the robustness of these DEGs and that *CADM2* RS1 deletion leads to broader transcriptomic changes. **b.** Example of functional enrichment of DEGs between one two-way comparison (hom 500 bp del and het 100 kb del). Enrichment is in terms related to neural development. Terms shown are those that are significant for both models individually and also for the convergence of these two DEG sets.

We performed pathway analyses on the iN DEGs (Fig. 4b, Table S11), focused on the most substantial perturbation across our study (hom 500 bp del) and the allele most closely matching patient variants (het 100 kb del). Hom 500 bp del DEGs (FDR <0.1) are enriched for terms including neurogenesis, neuron differentiation, and biological adhesion. Het 100 kb del DEGs are enriched for terms including cell migration, neurogenesis, neuron differentiation, and neuron development. We also performed gene ontology analysis of DEGs. The hom 500 bp del and het 100 kb del DEG sets – individually and the shared DEGs between them – were enriched for terms associated with neuronal growth and development including neurogenesis, neuron differentiation, and neuron development. While some genes with known ADHD-related genes were among the DEGs (e.g. *COMT* at FDR<0.1 in hom 500 bp del; *FOXP2*, *SNAP25*, and *ADRA2A* at nominal significance in het 100 kb del and hom 500 bp del), a human phenotype ontology (HPO) based analysis did not identify significant gene sets.

### Induced neuron models of CADM2 RS1 deletion do not show gross electrophysiological or morphological differences

CADM2 is involved in neurodevelopment via axon pathfinding/neurite outgrowth.^27–30^ To test for effects of *CADM2* RS1 deletion on gross neuronal morphology, we performed IncuCyte neuronal analysis on the cultured iNs. There were no differences among genotypes in neurite length per cell body cluster area (p=0.211, one-way ANOVA) or neurite branch points per cell body cluster area (p=0.406) (Fig. S10a-b), including at additional timepoints and cell plating densities (Fig. S11). To test for an effect of *CADM2* RS1 deletion on gross neuronal function, we performed microelectrode array analysis on the iNs and again found no significant functional impact on weighted mean firing rate (p=0.217, one-way ANOVA), average burst frequency (p=0.367), or other parameters (Figs. S10c-d, S12).

### A rodent Cadm2 RS1 deletion model demonstrates modest changes to functional brain connectivity and locomotor activity

*Cadm2* is known to undergo RS in the mouse brain;^7^ however, RS in rats has not been previously reported. We examined the rat genome and identified locus conservation, including an apparent RS1 site (Table S3). Rodents have been used previously as models of ADHD^31–33^ including to study *Cadm2*.^18^ We used CRISPR to ablate the *Cadm2* RS1 site in rats either locally (322 bp, “short” deletion) or on the scale of the deletions seen in patients (85 kb, “long” deletion), both heterozygously or homozygously (Fig. 5a). Deletion of RS1 by either the short or long allele ablated RS in cerebellum in a dose-dependent manner, as assessed by qPCR (Fig. 5b-c), paralleling our results from human iPSCs and iNs.

**Figure 5.**
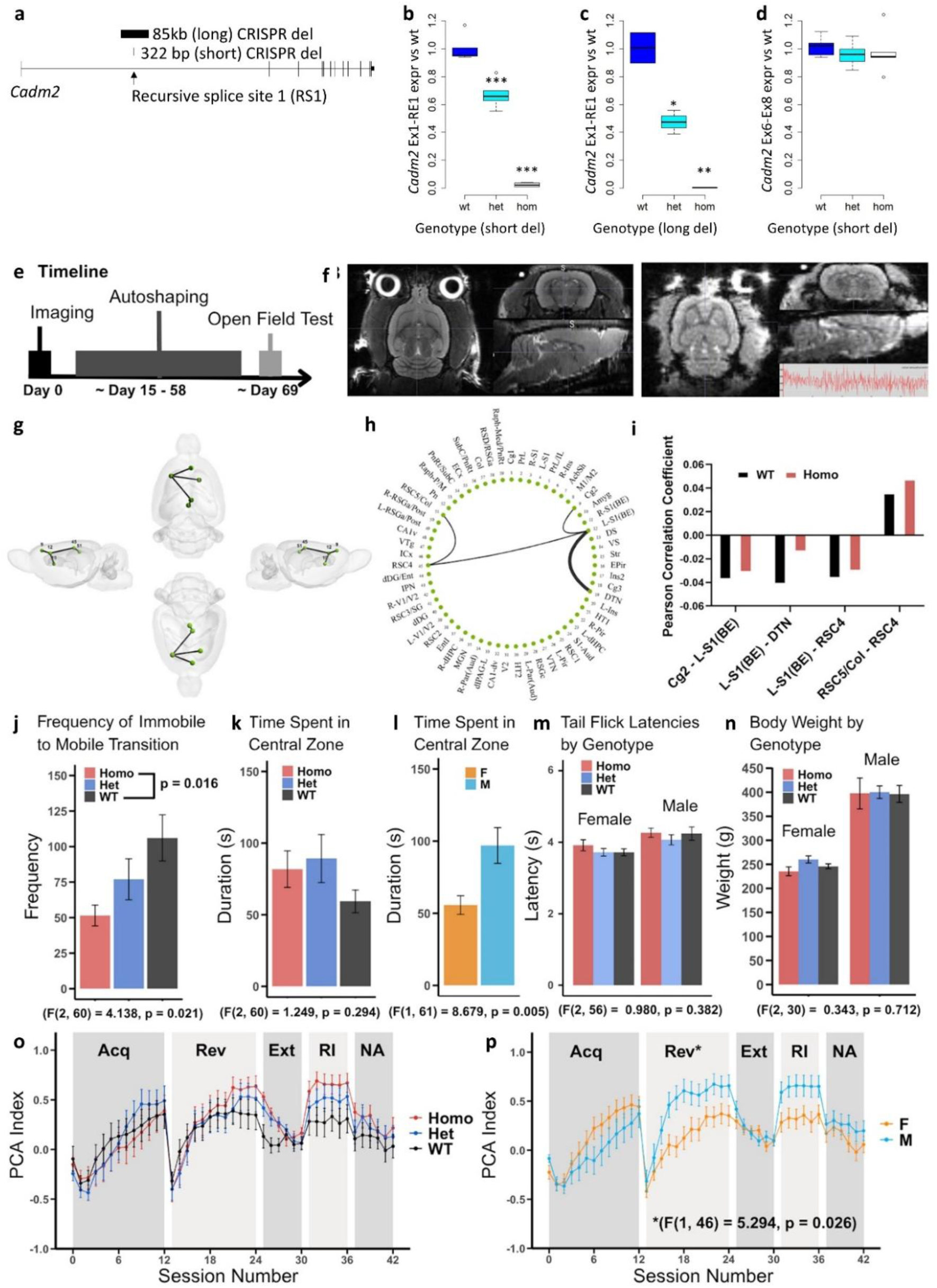
Changes to local gene expression, behavior, and functional brain MRI (fMRI) in rats following deletion of *Cadm2* RS1. **a.** Rat *Cadm2* locus. The genomic orientation of rat *Cadm2* is opposite to that of humans, but it is shown in the same orientation here for clarity. **b-c.** Recursive splicing is ablated in a zygosity dependent manner in (b) short deletion rats (110-112 days old; n=5 wt, 6 het, 6 hom) and (c) long deletion rats (74 days old; n=2 wt, 3 het, 2 hom from the same litter). Data are qPCR from exon 1 to RE1 of cerebellum-derived cDNA, normalized to *Polr2a*. For (b), one-way ANOVA p=1.51e-11, Tukey post-hoc HSD test het-wt FDR=1.25e-5, hom-wt FDR=0. For (c), one-way ANOVA p=0.0014, Tukey post-hoc HSD test het-wt FDR=0.0089, hom-wt FDR=0.0012. * p<0.05, ** p<0.005, *** p<0.0005. **d.** Deleting the *Cadm2* RS1 site does not significantly alter the expression of *Cadm2* in the cerebellum of short deletion rats (110-112 days old; n=5 wt, 6 het, 6 hom). Shown is cDNA qPCR from exon 6 to exon 8, normalized to *Polr2a*. One-way ANOVA p=0.695. **e-p.** fMRI, behavior, and body weight in long del rats. Extended long del behavior is in Fig. S17. Abbreviated short del behavioral analyses and body weight measurement were also performed (Fig. S18). **e.** Experimental timeline with approximate days spent in each phase, zeroed to date of first scan. **f.** Representative single-subject T2-weighted anatomical image (left) and one volume from an fMRI dataset (right). **g.** Three-dimensional rendering of the rat brain showing the subnetwork identified by Network-Based Statistic (NBS) analysis from multiple anatomical views. The subnetwork includes 4 edges and 5 nodes, reflecting significantly stronger functional connectivity in wt rats compared to hom long del rats (p = 0.019). Nodes are labeled by SIGMA atlas indices. **h.** Circular graph representation of the same significant subnetwork identified by NBS. Edges are visualized between region labels placed according to the SIGMA atlas. Edge thickness reflects the strength of the test statistic, with thicker lines indicating more significant connections. **i.** Groupwise average Pearson correlation coefficients for the edges identified by NBS. These connections exhibited significantly lower connectivity in hom long del rats. **j-l.** Open field test. **j.** Wt rats exhibited a significantly higher number of transitions from immobile to mobile compared to hom long del rats. **k.** Time in the central zone did not vary by genotype. **l.** Male rats spent a significantly greater cumulative duration in the center-zone compared to females (F(1, 61) = 8.679, p = 0.005) and made more transitions from center-zone to outer-zone (not shown; F(1, 61) = 4.323, p = 0.042). **m-n.** The tail-flick latency and body weights of age-matched rats did not differ significantly by genotype. **o-p.** Autoshaping results reveal no main effect of genotype on overall Pavlovian conditioned approach (PCA) Index (o) but show a main effect of sex (p), where males demonstrated greater sign-tracking tendencies than females during the reversal stage of the task (F(1, 46) = 5.294, p = 0.026). Abbreviations: Cg2 – Cingulate Cortex area 2; L-S1(BE) – Left Primary Somatosensory Cortex (Barrel Field); DTN – Dorsal Thalamic Nucleus; RSC4 – Retrosplenial Cortex area 4; RSC5/Col – Retrosplenial Cortex area 5 / Colliculus; Acq = Acquisition; Rev = Reversal; Ext = Extinction; RI = Reinstatement; NA = Negative Automaintenance.

To assess for differences in the brains of living rats, functional MRI (fMRI) data were acquired from 58 rats that were hom, het, and wt for the long del (Fig. 5e). Following quality control, 3 datasets were excluded due to motion-related or slice-timing artifacts, resulting in final images from 54 rats for analysis (Wt: 6M, 11F; Het: 10M, 10F; Hom: 10M, 7F). Representative anatomical and functional images are shown in Fig. 5f. Canonical resting-state networks commonly reported in rodents were preserved across subjects. Network-Based Statistic (NBS) analysis identified a significant subnetwork (p=0.019, by permutation-based network component testing) composed of 4 edges and 5 nodes with decreased connectivity in hom long del rats relative to wt (Fig. 5g-h). This network included connections between the cingulate cortex (Cg2) and the left primary somatosensory cortex, as well as between the somatosensory cortex and both the dorsal thalamic nucleus and retrosplenial cortex (RSC4). An additional connection was found between RSC4 and RSC5/colliculus. Test statistics for these connections ranged from 3.12 to 3.66, surpassing the pre-defined threshold of 3. Groupwise correlations for these connections are illustrated in Fig. 5i.

*Cadm2* has been demonstrated to regulate body weight and activity in mice and BMI in humans.^18,34^ The body weights of age-matched rats did not differ significantly by genotype among long del (F(2,30) = 0.343, p = 0.712, Fig. 5n) or short del (F(2,41) = 1.865, p = 0.168; Fig. S18f) cohorts. Of note, the aforementioned mouse models involved complete deletion of *Cadm2*, and human BMI association involved GWAS SNPs, whereas in our rat models only the intronic RS1 site was deleted.

We tested for ADHD-like behaviors in long del rats, as well as broader behavioral changes including reinforcement and reversal learning, and risk taking (Fig. 5j-p). On the open field test, we observed a main effect of genotype (F(2, 60) = 4.199, p = 0.020), with hom long deletion rats exhibiting significantly fewer transitions from immobile to mobile compared to wt (p = 0.016; Fig. 5j). When animals were assessed for Pavlovian conditioned approach (PCA) (sign-tracking vs. goal-tracking), which has previously been tested in humans and associated with externalizing behaviors, there was no main effect of genotype on PCA Index (Fig. 5o). There was a main effect of genotype on breakpoints in the progressive ratio breakpoint test (PRBT) ([F(2,33)=3.480, p=0.043]; Fig. S17q), which revealed het long del rats tended to have lower breakpoints compared to both wt rats (*p*=0.096) and hom long del rats (*p*<0.05). We assessed pre-pulse inhibition (PPI) on hom and het long del and wt rats, with no main or interactive effects of genotype across each measure (Fig. S17a-c). There were no long del genotype differences observed on any of the primary or secondary outcome measures of the Iowa Gambling Task (IGT) (Fig. S17d-e), except for the number of choice trials completed ([F(2,31)=4.445, p=0.020]; Fig. S17e). Group comparisons revealed that het long del rats completed more trials in the IGT than wt rats (*p*<0.01) and tended to complete more trials than hom long del rats (*p*=0.069; Fig. S17e). There were no long del genotype differences observed on any exploratory measures (Fig. S17f-k), or reversal learning or reinforcement learning from probabilistic reversal learning (PRLT) measures (Figure S17l-p).

Additional screens in short del rats identified differences in mutant animals (Fig. S18). Maximum time spent mobile was greater in hom rats compared with het (p=0.028 by a 2-way ANOVA test; Fig. S18b). Body elongation percentage was greater in het rats than wt (p=0.036 by a 2-way ANOVA test; Fig. S18c). Tail flick pain response (p=0.021 by a 2-way ANOVA test; Fig. S18d) and hot plate pain response (p=0.029 by a 2-way ANOVA test; Fig. S18e) were decreased in hom vs het animals.

Because the above behavior differences were modest and/or inconsistent between short and long del trials, we wondered whether the expression of *Cadm2* is less reliant on recursive splicing in rat than in humans. qPCR of short del cerebellums identified no significant change in *Cadm2* expression upon deletion or RS1.

## Discussion

Building upon our discovery of a strong association between deletions of an RS site in *CADM2* and neurobehavioral disease risk,^11^ we characterized the population allele spectrum, molecular mechanisms, and cellular and phenotypic consequences of these deletions in relevant *in vitro* and *in vivo* systems. SV detection from WGS data in gnomAD and All of Us allowed us to characterize deletion alleles in the general population. Human induced neurons engineered with *CADM2* RS1 deletions or inversion exhibited: 1) ablation of recursive splicing; 2) altered pattern of RNA content across *CADM2*’s first intron, 3) decreased overall *CADM2* expression; 4) alterations to *CADM2* transcript usage; and 5) global differentially expressed genes enriched for neuronal processes. These findings demonstrate functional roles for RS in a human gene. Rats engineered with *Cadm2* RS1 deletions showed similar effects on RS usage, as well as modest behavioral differences and altered functional brain connectivity between regions previously implicated in ADHD.^35,36^

### The purpose of recursive splicing

The purpose of recursive splicing has not been entirely clear; transcript choice and splicing fidelity across long introns have each been suggested. Sibley et al.^7^ targeted the zebrafish *cadm2a* RS exon’s 5′ss with an antisense oligonucleotide (ASO) and showed decreased expression of downstream exons, but a similar experiment for the human *CADM1* and *ANK3* genes in a neuroblastoma cell line did not demonstrate this effect. Furthermore, there was no significant change in splicing from exon 1 to 2 for zebrafish *cadm2a*. Limitations of that study included the use of only qPCR, different genes being tested in different species, and the use of ASOs rather than deleting the RS sites.

Here, we show that deletion of the RS1 site in human iPSCs and iNs decreases overall *CADM2* expression and affects transcript representation of this gene. The decrease of all robustly expressed exons downstream of RE1, but not of the exon preceding it (exon 1), upon ablating recursive splicing suggests a lower efficiency of the spliceosome joining exon 1 to the next constitutive exon, potentially owing to the extreme distance between them (e.g. 767 kb from exon 1 to 2) and/or differing dynamics of a pre-mRNA with the 279 kb exon-1-to-RS1 segment retained. Additionally, our data show that RE1 and RE2 are included in *de novo* reconstructed transcripts and that RE1 can splice to RE2. This latter observation suggests that RE1’s native 5′ss can be used, i.e. RE1 is not always recursive. It remains unknown the extent to which transcripts containing RE1 and/or RE2 are intermediates from which one or both of these recursive exons will be subsequently removed – either sequentially or in a single step – or whether they are fully spliced but non-productive transcripts that will be degraded by NMD because of stop codons in the REs. In either case, their inclusion may play a role in modulating mRNA and/or protein levels; for example, deletion of RE1 may increase the production of CADM2+RE2 (Fig. 3c) at the expense of productive splicing to exon 2. Of note, splicing of neither RE1 to RE2, nor of RE2 to exon 2, is expected to reconstitute a strong 5′ss (Figs. S5, S6); thus, once an RE’s native 5′ss used, it appears unlikely that the RE would subsequently be spliced out.

In human iPSCs and iNs, we modeled larger (mimicking patient/population alleles) and smaller (specifically ablating only RE1) alleles. While there was evidence of a dosage (e.g., zygosity) effect, there was no consistent evidence that the larger and smaller deletions behaved differently with regard to their effect on local or transcriptome-wide gene expression. This, and similar effects with inversion (without sequence loss of RE1), suggest that the phenotypic effect of the patient/population alleles may result primarily from disruption of recursive splicing. We also modeled both larger and smaller deletions in rats. While there were some disparate results depending on the size of the deletion, these experiments were not performed as a single batch containing large and small del animals, potentially limiting the ability to distinguish differential effects based on deletion size.

While human RS was initially discovered in the brain^7^ where many very long genes are expressed and where exon junction complex components that suppress RS are reportedly more lowly expressed,^13,14,37^ RS has been identified in other tissues.^4,38,39^ We show RS is active in both human iPSCs and induced glutamatergic neurons.

### Phenotypic associations with variation in CADM2

The *CADM2* locus houses copious GWAS and PheWAS hits for numerous psychobehavioral traits related to ADHD (e.g., impulsivity, substance use) and beyond (e.g., educational attainment, BMI) (https://www.ebi.ac.uk/gwas/, https://disgenet.com/, https://platform.opentargets.org/, ^16–18,40–42^. Only a few studies have investigated functional effects of *CADM2* variation: a subset of GWAS SNPs colocalize with *CADM2* eQTLs,^17,43^ and some GWAS SNPs increase *CADM2* expression in human hypothalamus.^34,44^ Supporting the pleiotropic effects of variation at this locus suggested by GWAS, *Cadm2*-deficient mice not only display differences in impulsivity-related traits (e.g., less risky choices on the Iowa gambling task), but also exhibit increased locomotion, energy expenditure rate, and core body temperature.^18,34^ Although not proof of causality, neural stem cells and neurons from bipolar disorder patients exhibit differences in CADM2 protein expression.^45^ Intriguingly, while some GWAS and PheWAS SNPs localize near *CADM2* RS1, the gene’s major hotspot is the minor promoter/RS2 site (https://www.ebi.ac.uk/gwas; ^41^) (Fig. S15). This region has a substantial H2K27Ac peak and is the origin of a 3D architectural stripe connecting back to the major promoter (UCSC Genome Browser; not shown). This suggests important future studies of both rare and common alleles in that region of the gene.

It will be interesting to learn the full breadth of phenotypes associated with *CADM2* RS1 variation. Additional large cohorts with WGS data will be required to replicate the association of *CADM2* RS1 deletions with ADHD and the phenotypes nominally associated with RS1 deletions in our discovery efforts (bipolar affective disorder, abnormality of movement, and impairment in personality functioning), and to discover novel associations. In other words, *CADM2* RS1 variation may constitute a pleiotropic risk factor for several pathophysiologically-related neuropsychiatric traits.

### Limitations and future directions

Because RS1 deletion is rare, we have not had an opportunity to deeply phenotype the neuropsychiatric profiles of individuals with this mutation. Our use of an animal model is intended to partially address this limitation; however, the behavioral phenotypes observable in rodents were modest and present their own limitations regarding relevance to specific human traits such as a diagnosis of ADHD, particularly in light of our finding that rat *Cadm2* expression (in cerebellum for short del animals, at least) is apparently not as dependent on RS1 as it is in human iPSCs and iNs. RS has been identified in additional genes including several associated with neurodevelopmental disorders (Table S2; ^46^). To preliminarily identify whether these RS sites are subject to deletion or mutation at the population level, we investigated gnomAD (v4.1.0). RS site deletions not also affecting a coding exon were found for 5 of 10 RS sites, all of which were rare (allele frequency ≤3.17e-5) (Table S9). SNVs in a 10 bp window including each RS site were found for 7 of 10 loci (Table S9), also all rare (allele frequency 1.91e-4 to 1.02e-6). Thus, variation potentially affecting RS in other loci known to use RS exists in the population, opening the possibility of additional phenotype-associated variation altering RS at these loci. Additional questions opened by our work include the role of RS in additional cell types and developmental timepoints (Fig. S7), as well as experimental/therapeutic modulation of splicing or expression in *CADM2*.

## Acknowledgments

The authors thank Riya Bhavsar, Zsabre Wright, Laura Smith, Lynn Malloy, Drs. Gabi Xavier, Bimal Jana, Colby Chiang, Madelyn Light, and Kyle Satterstrom for assistance with and helpful discussions about the project. This work was supported by the National Institutes of Health (NHGRI U01HG011755, NINDS K08NS117891, NIMH R01MH115957, R01MH123155, NICHD P50HD104224), the National Institute on Drug Abuse (NIDA; P50DA037844, P50DA054071, DP1DA054394, and P30DA060810), the Boston Children’s Hospital (BCH) Office of Faculty Development, the BCH Basic and Clinical Translational Research Executive Committee, the BCH Tommy Fuss Center for Neuropsychiatric Disease Research, the Massachusetts General Hospital Executive Committee on Research Fund for Medical Discovery, the California Tobacco-Related Disease Research Program T32IR5226, Canada Research Chair-CIHR (JYK).

## Declaration of Interests

The authors declare no competing interests.

## Declaration of generative AI and AI-assisted technologies

AI was utilized to generate the rat silhouettes in the graphical abstract.

## List of Supplemental Tables

Table S1. CRISPR cell line genotypes and names.

Table S2. Recursive splice sites from Sibley et al Nature 2015.

Table S3. Coordinates of *CADM2* landmarks.

Table S4. *CADM2* exon definitions.

Table S5. Probes for capture RNA sequencing.

Table S6. Primers, CRISPR guides, ddPCR assay.

Table S7. Rat genotypes, crosses, IDs.

Table S8. Associations with HPO terms (metaphenotypes) in rCNV analysis.

Table S9. gnomAD and All of Us alleles affecting RS sites.

Table S10. Differentially expressed genes (DEGs).

Table S11. SynGo and GSEA functional pathway enrichment data.

Table S12. Exon expression statistics.

Table S13. Transcript Quantification.

## List of Supplemental Figures

Figure S1. Example genotyping of *CADM2* RS1 deletions and inversion in CRISPR engineered iPSCs and gene-edited rats.

Figure S2. Example induced neuron morphology. Figure S3. RNA-seq quality control.

Figure S4. Sashimi plots.

Figure S5. *CADM2* isoform census in human brain RNA.

Figure S6. *CADM2* RE1 and RE2, as defined by total RNA-seq in human iNs.

Figure S7. Differences in *CADM2* expression between iPSCs and iNs.

Figure S8. Exon, junction, and transcript expression, extended.

Figure S9. CRISPR *CADM2* RS1 site deletion in human induced pluripotent stem cells (iPSCs) ablates recursive splicing, alters the pattern of intronic mRNA abundance, and decreases *CADM2* expression.

Figure S10. *CADM2* RS1 deletion does not alter gross neurite morphology or neuronal activity.

Figure S11. Extended IncuCyte live cell analysis data.

Figure S12. Extended microelectrode array data.

Figure S13. Additional junction expression analyses, iNs.

Figure S14. *CADM2* RS1 deletions and simple nucleotide variants (SNVs) in gnomAD.

Figure S15. GWAS hits in *CADM2* are overrepresented in the region of a minor promoter and the second recursive splice site (RS2).

Figure S16. Smoothened read depth over *CADM2* in iNs, by genotype.

Figure S17. Extended long del rat behavioral data.

Figure S18. Behavior and weight in *Cadm2* short del rats.

Figure S19. Fold change of *de novo* reconstructed transcripts in iNs, extended.

## List of Supplemental Files

De novo reconstructed transcripts.bed

## Methods

### Additional cohort analyses

For deletions overlapping the *CADM2* RS1 site in gnomAD-SV v4 (https://gnomad.broadinstitute.org/gene/ENSG00000175161?dataset=gnomad_sv_r4), we performed additional quality control by manually assessing the sequencing depth profile of large copy number variants. To determine if there were any SNVs or indels linked to the gnomAD deletions affecting *CADM2* RS1, we computed linkage disequilibrium between each deletion and all nearby short variants within 1 Mb. Linkage disequilibrium was calculated across all gnomAD samples with both SNV/indel and SV calls (N_ALL_ = 62,877) as well as only those samples in the same genetic ancestry group as each RS1 deletion (N_EAS_ = 2,012; N_NFE_ = 29,488; N_SAS_ = 2,185).

### Induced pluripotent stem cell (iPSC) culture

Cell culture was performed under MassGeneralBrigham biosafety protocol 2011B000500. Wildtype human iPSCs, designated MGH2069, were a gift from S. Haggarty. This line was originally derived from fibroblasts of a healthy adult female.^47^ No artifactual variants of interest were identified via array comparative genomic hybridization at P27/28 or whole genome sequencing (WGS) at P30 (data available upon request). Mycoplasma was absent by PCR (data not shown). Cells were cultured on Matrigel (Corning) with Essential 8 (E8) medium (Gibco) with 1% penicillin-streptomycin (Fisher Scientific), passaged using ReleSR (STEMCELL Technologies), cryopreserved with mFreSR (STEMCELL Technologies), and CRISPR edited at P35. Y-27632 ROCK inhibitor (Biological Industries) was supplemented at 10uM for 24h after thawing or passaging, and during FACS.

### CRISPR mutagenesis

Guide RNAs for dual-guide CRISPR (see ^48^) were designed using the UCSC Genome Browser CRISPR track and the IDT guide RNA design checker (https://www.idtdna.com/site/order/designtool/index/CRISPR_SEQUENCE) and ordered as sgRNAs from IDT. Guide sequences are listed in Table S6. For human cell CRISPR mutagenesis, two different yet closely positioned pairs of guides were designed per desired allele but utilized as independent pairs. Human iPSCs were passaged at least twice after thawing prior to transfection, including 24h before. CRISPR components were based on the Alt-R CRISPR-Cas9 system (IDT) and delivered as in (http://sfvideo.blob.core.windows.net/sitefinity/docs/default-source/user-guide-manual/alt-r-crispr-cas9-user-guide-ribonucleoprotein-transfections-recommended.pdf?sfvrsn=1c43407_8), with modifications. Briefly, Alt-R S.p. HiFi Cas9 nuclease V3 (IDT) and sgRNA guide pairs were diluted in duplex buffer (IDT) and mixed at RT x 5 min to form ribonuclear proteins (RNPs). RNPs were co-transfected with CleanCap EGFP mRNA (TriLink Biotechnologies) into ∼50% confluent iPSCs using Lipofectamine STEM (Thermo-Fisher) diluted in Opti-MEM I (Gibco). Media was changed to E8 at 24h and cells were harvested for FACS at 48h. Additional details (e.g. dilution volumes) are available on request.

### FACS sorting, genotyping, and pluripotency selection

Edited iPSCs were FACS sorted using the FACSAria system (BD), gating on the highest 10% GFP-expressing live, single cells. Sorting enabled clonal wt and edited lines to be obtained. Genotyping was performed via a combination of PCR and ddPCR and repeated to ensure accuracy. PCR utilized 2X PCR Master Mix (Promega) and the following program: 95C x 2 min; 37 or 40 cycles of 95C x 30 sec, 57C x 30 sec, 72C x 2 min; 72C x 5 min. ddPCR was done with Bio-Rad reagents and machines including HEX control probe Assay ID dHsaCP2500350, Automated Droplet Generator, QX200 Droplet Reader, and QuantaSoft software, except for a Thermo-Fisher Taq-Man target probe (ID CADM2rs1_CDYMJ4M; proprietary sequence). AluI or HindIII enzymes were utilized if digestion was done. The ddPCR program was: 95C x 10 min; 40 cycles of 94C x 30 sec, 57C x 1 min; 98C x 10 min. PCR primers and the ddPCR amplicon region are listed in Table S6. Sample genotyping results are in Fig. S1.

Edited lines were thawed as P40-41. Four clones per genotype were carried forward for further experimentation, except for inversions and homozygous deletions, where lines were duplicated or quadruplicated, respectively, to derive four pseudo-biological replicates (Table S1). Before starting iN differentiation, edited iPSCs underwent magnetic column-based selection (MACS; Miltenyi) for pluripotency based on TRA-1-60 positivity.

### Induced neuronal (iN) differentiation

Edited, pluripotency-selected iPSCs were thawed onto Geltrex (Gibco) simultaneously as P41-2. iPSCs were harvested one passage later in TRIzol (Thermo Fisher). Differentiation to induced glutamatergic neurons (iNs) was carried out using a 24-day doxycycline responsive lentiviral-mediated *Ngn2* transgene approach.^49,50^ Milestones of the protocol include: infection with rtTA– and pTetO-mNgn2-encoding lentiviruses in the presence of polybrene (Sigma Aldrich) on day –8; culture with Neuronal Maintenance Medium (Thermo Fisher) on days 0-24; passaging with Accutase (Thermo Fisher); *Ngn2*-induction with doxycycline (Sigma Aldrich) on days 0-10; selection with puromycin (Thermo Fisher) on days 1-3; plating onto poly-L-ornithine (P-L-O; Sigma-Aldrich) laminin (Sigma) plates on day 4; treatment with Ara-C (Sigma) on day 5; and addition of NT-3 (Preprotech) and BDNF (Prospec Bio) on days 4-24. Modifications included the use of 0.1% laminin. Differentiation was monitored via inspection of cell morphology (Fig. S2), RNA expression of marker genes (Fig. S3), and by performance on imaging and electrophysiologic assays (see below). Neurons were harvested in TRIzol on day 24.

### RNA isolation

TRIzol samples underwent standard chloroform extraction, isopropanol-based RNA precipitation, and 75% ethanol washing, with modifications including the use of Phasemaker tubes (Thermo Fisher) and RNAse-free glycogen (Roche). RNA was quantitated using a Nanodrop spectrophotometer (Thermo) and QC’ed using a TapeStation 4200 (Agilent). To obtain an initial census of *CADM2* transcripts and recursive splicing in wt humans and rats, human whole brain RNA and rat brain total RNA were purchased from Clontech (TaKaRa).

### Library preparation and sequencing

Total RNA-seq libraries were prepared with the NEBNext rRNA Depletion Kit (New England Biolabs) following the manufacturer’s instructions. In brief, RNA samples were hybridized with rRNA depletion probes, then treated with RNase H and DNase I to deplete ribosomal RNAs. RNA molecules were fragmented using divalent cations and higher temperature, followed by first-strand cDNA synthesis primed with random primers. Strand specificity was achieved by replacing dTTP with dUTP during second-strand cDNA synthesis. cDNA libraries were end-repaired and A-tailed, followed by universal hairpin loop adapter ligation and USER enzyme digestion. Libraries were indexed via an 11-cycle PCR amplification using sample-specific dual-indexed primer pairs. QC of the prepared libraries’ size distribution and concentration was undertaken on a TapeStation 4200 (Agilent) and a qPCR with a Library Quantification Kit (Kapa Biosystems), respectively. Finally, libraries were pooled in equimolar amounts, then sequenced on an Illumina NovaSeq6000 using an S4 300 cycles kit.

For targeted capture (Cap-seq), total RNA-seq libraries were multiplexed into batches of 12 (up to 8 ug of total material). Targeted enrichment was performed using custom capture baits from IDT (see below) via the IDT xGen Hybridization Protocol. In brief, multiplexed libraries were combined with Human Cot DNA and xGen blocking oligos and dehydrated prior to resuspension in hybridization buffer and baits. After four hours of incubation, bait-hybridized libraries were combined with buffer-resuspended streptavidin beads and several washes performed to remove any non-hybridized libraries, followed by 15 rounds of on-bead, post-capture PCR. PCR amplified libraries were purified using SPRI bead clean up and enriched libraries were analyzed via TapeStation and Qubit. Finally, libraries were pooled in equimolar amounts, then sequenced on an Illumina NovaSeq6000 using an S4 300 cycles kit.

### Capture reagent

A capture reagent was designed in conjunction with IDT. Probes (Table S5) were chosen from among the IDT Discovery Pool set, with high and medium risk probes removed. Exons and other sequences of interest within *CADM2* were padded by 100 bp for the design, with the exception of the alternative first exon being padded by 500 bp and the recursive exons by 1000 bp.

### qPCR and RT-PCR

cDNA was prepared using SuperScript IV reverse transcriptase (Thermo Fisher), with both oligo (dT)_15_ (Promega) and random primers (Promega) in order to represent both mature and pre-mRNA species (particularly in light of *CADM2’*s extremely long annotated 3′ UTR), according to the SuperScript protocol (https://assets.thermofisher.com/TFS-Assets%2FLSG%2Fmanuals%2FSSIV_First_Strand_Synthesis_System_UG.pdf). qPCR was performed in technical duplicate with SYBR Green I Master mix (Roche) on a LightCycler 480 instrument (Roche), with a dilution series for each experiment to calculate reaction efficiency, and utilizing the following program: 95°C x 5 min (ramp rate 4.4°C/sec); then 35 cycles of 95°C x 10 sec (ramp rate 4.4), 55C x 20 sec (ramp rate 2.2), and 72°C x 30 sec (ramp rate 4.4). RT-PCR utilized the same reagents and program as genotyping PCRs (above). Sanger sequencing of RT-PCR products was performed at the Massachusetts General Hospital DNA Core. qPCR and RT-PCR primers are listed in Table S6. Some primer sequences were obtained from published reports.^7,51,52^

### RNA-seq processing

Total RNA-seq reads were aligned to the Ensembl Human Genome Reference GRCh38.92 using STAR v2.5.2b allowing a 5% mismatch rate.^53^ Trimmomatic (v0.36) was used to trim Illumina TruSeq adapters to a minimal length of 105 bp (2:30:10 LEADING:3 TRAILING:3 SLIDINGWINDOW:4:20 MINLEN:105).^54^

### Establishing RS pattern

Recursive splicing (RS) is characterized by an exon-like sawtooth pattern of RNA abundance. To establish this RS pattern in the first intron of the *CADM2* gene containing a recursive splice site (RS1), we calculated the slope of coverage of the first intron. The sawtooth pattern was defined by dividing the first intron into three segments (first: end of exon 1 (chr3:84959669) to RE1 start (chr3:85238956), middle: RE1 (chr3:85238957) to RS2 exon start (chr3:85511857), end: RS2 exon (chr3:85511858) to exon 2 start (chr3:85726144)). Total RNA-seq read counts were normalized per sample, and for each position in the first intron the positions were converted to relative positions by subtracting the position by the start position of the segment and scaling by 1 million. The slope for each segment in each sample was calculated by fitting a linear model using the normalized read count to the relative position. Slopes were then used to compare the effect of genotypes of the saw tooth pattern. Significance was determined using a t-test.

### Differential gene expression analysis

HTSeq (v 0.11.2) was used to perform gene counts on uniquely mapped reads obtained from total RNA-seq.^55^ To ensure compatibility with the Illumina TruSeq libraries, the “-s reverse” option was used to indicate strandedness and only unique reads were counted. Differential expression analysis was performed by comparing unedited samples (wt) to CRISPR-edited genotypes. Genes were first filtered to only allow median expression of ≥0.1 counts-per-million in either wt or edited samples. These genes were kept for downstream analysis in the Bioconductor package DESeq2 (v 1.34.0).^56^ Raw gene counts were then normalized using sample-wise size factors. Y chromosome genes were removed from the analysis as the cell lines did not have Y chromosomes and any mapping to these regions would be artefacts and arise from pseudoautosomal regions or highly similar regions elsewhere in the genome. One inversion iN sample (rC5_2) was removed from analyses as it was a *CADM2* expression outlier.

### Genome guided de novo transcript reconstruction and recalibration

To perform genome-guided *de novo* assembly, total RNA-seq and Cap-seq alignments within the *CADM2* gene region +/− 1 kilobases were extracted using samtools,^57^ duplicate reads were removed and read pairs from the same genotype in iNs were merged. Transcripts were assembled using Trinity v2.20 for each genotype (wt, hom 500 bp del, het 100 kb del, het 500 bp del, het 500 bp inv) with the following parameters “-SS_lib_type RF” and “-genome_guided_max_intron 100000”.^58^ Assembled transcripts were compared to chromosome 3 from human genome GRCh38.92 using BLAT and selected to be mapped within the CADM2 gene region, resulting in 48663 transcripts across the five genotypes. These *CADM2* region transcripts from iNs were merged into a non-redundant set based on 95% similarity. 1182 transcripts of these pass junction TPM cut-off > 1e-6 and further merging assembled transcripts that were fragments of the same transcripts based on their exon boundaries, and resolving internal splicing structure, resulted in 1173 transcripts. Merging within genotypes and then across genotypes to combine transcripts with a similar structure yielded 891 transcripts. Seventeen unique transcripts had support in more than 50% of samples of any genotype. All transcripts were annotated according to their structures matching the GRCh38.p14 Ensembl (v112) *CADM2* transcripts and recursive exons were based on published sources.^4,7^

Expression of these transcripts was then quantified using RSEM v1.1 against a reference built against the *de novo* transcripts.^59^ We used transcript-per-million (TPM) and absolute counts to determine the expression profiles of the transcripts and identified significantly different transcripts and fold change via a t-test using wt samples as a baseline.

### Differential Exon Expression

*DEXSeq*^60^ was used to assess differentially expressed exons in *CADM2* using exon definitions as in Table S4. HTSeq was used to count the number of total RNA-seq reads mapping to each exon and *DEXSeq* was used to model exon expression using a generalized linear model on negative binomial distribution. The model takes biological variation into account to estimate dispersion and size factors, fit the dispersion-mean relation and test for differential exon usage. *DEXSeq* calculates p-values based on the Fisher’s exact test with high sensitivity concerning the wt lines. The p-values are corrected following the Benjamini-Hochberg procedure. We used read mapping to *CADM2* to estimate the exon expression in each mutant genotype, compared to wt, by applying a t-test to the counts.

### Gene Ontology Analysis

To identify enriched molecular signatures in the edited lines, the Gene Set Enrichment Analysis GO resource was used (https://www.gsea-msigdb.org/, v. MS1). A statistical threshold of FDR < 0.1 was used. Fisher’s Exact test was used to test for the enrichment of GO terms. Adjusted p-values are calculated using the Benjamini-Hochberg procedure.

### Microelectrode array analysis (MEA)

*Ngn2*-infected iPSCs were thawed at P31, then differentiated to iNs as above. On day 4, cells were plated on PEI (polyethylenimine; Sigma-Aldrich) coated 48-well CytoView MEA plates (Axion Biosystems) in 1.5% laminin (Sigma). At least 6 wells per line/replicate (24 wells per genotype) were plated, alternated by genotype to limit plate edge effects. Media was changed to BrainPhys Neuronal Medium (STEMCELL Technologies) on day 24. Measurements on the Maestro Pro (Axion) instrument were made for 15 minutes every three days using “Neural Real-Time” standard settings in Axis Navigator and without supplemental CO_2_. Data were analyzed using AxIS Metric Plotting Tool software with default settings. Day 27 was chosen for analysis, being the differentiation day with the highest weighted mean firing rate and the highest number of genotypes represented by wells with ≥ 8 active electrodes.

### IncuCyte automated neuronal analysis

*Ngn2*-infected iPSCs were thawed, then differentiated to iNs as above. On day 4, cells were plated on 96-well BioCoat P-L-O/Laminin plates (Corning). 3 wells per line/replicate (12 wells per genotype) were plated, alternated by genotype and avoiding edge wells to limit plate edge effects. Replicate plates featured either 10k or 15k cells/well. Live-cell morphological analyses were performed stain-free on the IncuCyte S3 instrument (Sartorius) in the Boston Children’s Hospital Human Neuron Core, per the following protocol: https://www.sartorius.com/en/products/live-cell-imaging-analysis/live-cell-analysis-instruments/s3-live-cell-analysis-instrument. Images were obtained every 4 hours on differentiation days 11-13 (time point 1), 18-20 (time point 2), and 25-27 (time point 3), resulting in 13 sets of images per time point. Observed values were averaged across the 13 measurements per time point. Images were analyzed via Neurotrack software (Sartorius) on default settings for branch points, neurite length, cell body clusters, and cell body cluster area.

### Rat models

CRISPR-Cas9 editing with single or dual sgRNAs flanking the target region was used to generate founder rats with the *Cadm2* RS1 site deleted, following the general protocol described in.^61^ The background is an outbred population of Heterogenous Stock (HS) rats.^62^. This background is an admixture of eight founder inbred strains and is intended to normalize for strain-specific effects. To maintain background heterogeneity, founder mutant animals and each subsequent generation were backcrossed to HS animals from non-sib pairs until heterozygous animals from different litters were selected for intercrosses to generate homozygous animals. Relevant sgRNA and primer sequences and the coordinates of the deletions and other *Cadm2* landmarks in rat are in Tables S3 and S6. Sample genotyping is in Fig. S1. Genotypes and IDs of founders and animals harvested for tissue collection are in Table S7.

Rat brains were snap frozen in liquid nitrogen. For expression analyses, cerebellums were sampled from frozen brains, ground in TRIzol, and RNA was extracted with an extra centrifugation to remove lipids. cDNA was prepared as above. Technical duplicates were inherent to qPCR. Additional technical duplication at the cDNA preparation step was performed as mentioned in the text.

### Magnetic resonance imaging

As in prior studies, rats were initially anesthetized in an induction chamber using 4–5% isoflurane with an oxygen flow of 1–1.5 L/min. Anesthesia was maintained via a nose cone at 2.0– 2.5% isoflurane with the same oxygen flow rate, and animals received an intraperitoneal injection of dexmedetomidine at a dose of 0.018 mg/kg. After positioning the animals in the MRI scanner, a continuous infusion of dexmedetomidine (0.018 mg/kg/h) was administered for the remainder of the imaging session. Isoflurane was gradually reduced from 2.0–2.5% to 0.8–1.0% over 15 minutes after initiating the dexmedetomidine infusion, maintaining the oxygen flow between 1–1.5 L/min to stabilize physiological parameters (respiration: 74 ± 5.2 breaths/min; heart rate: 314 ± 23 bpm). Body temperature was maintained at 37.0 ± 0.5 °C using an air-based heating system. EPI data acquisition commenced once physiological conditions were deemed optimal, consistent with previously published protocols.^63,64^

Images were acquired using a 9.4 T Bruker small animal MRI scanner at the Centre for Functional and Metabolic Mapping located within Robarts Research Institute at Western University.

T2 Anatomical images were acquired for each subject at the beginning of each session with T2-weighted TurboRARE pulse sequence (8 averages, 35 slices, slice thickness = 400 μm, FOV 38.4 × 24 mm, matrix size 192 × 120, in-plane resolution = 200 × 200 μm, TE = 39.0 ms, TR = 7.0 s, Echo Spacing = 11.00 ms, Rare Factor 8, total acquisition time = 14 min).^65^

Resting-state fMRI images were acquired based on optimized scan parameters,^66^ using a gradient-echo EPI sequence (4 runs, 400 volumes per run, TE = 15.0 ms, TR = 1.5 s, FOV 38.4 × 38.4 mm, matrix size 96 × 96, 35 slices, isotropic resolution = 400 μm, bandwidth 280 kHz). Pre-processing was conducted using RABIES software and the SIGMA rat brain template as one previously.^67–69^

### Statistical analysis (fMRI)

To examine differences in functional connectivity between wild-type and CADM2 mutant rats, the Network-Based Statistic (NBS) method was used.^70^. This approach is designed to detect statistically significant alterations in subnetworks and has been increasingly adopted for studies aimed at cross-species translational insights,^71^ which reflects a key objective of the present research. Unthresholded correlation matrices were utilized in the analysis to retain the full spectrum of connectivity values and to prevent potential biases associated with thresholding procedures.^70^ Group comparisons between wild-type and RS1 het and homo rats were conducted using two-tailed t-tests (p = 0.05). Subnetworks showing significant group-level differences were subsequently extracted and examined for interpretation.

### Open Field Test

Locomotor activity was measured using an open field test in Plexiglas cages (50 × 50 × 40 cm) with a gray floor and black walls.^72^ It was recorded via a GigE camera mounted 190 cm from the floor, and a PC running video tracking software (Ethovision v16.0.1536, Noldus, Wageningen, NL). After an initial habituation period to the testing room, Homo (n = 20), Het (n = 20), and WT (n = 23) rats were placed in the center of the arena and allowed to freely explore for 10 minutes. Parameters such as total distance traveled (cm), velocity (cm/s), movement duration, time spent in the center zone (the virtual area with a size of 25 × 25 cm located in the middle of each cage) were among the variables collected.

### Autoshaping

Sign-tracking behaviors, also known as Pavlovian Conditioned Approach (PCA) responses, have been classically studied through an autoshaping procedure, as previously described.^73,74^ Homozygous (Hom, n = 17), Heterozygous (Het, n = 18), and Wild-Type (WT, n = 17) rats underwent (a) magazine training until consistently eating sucrose pellet rewards (45 mg, Bio-Serv, NJ) (one to two days), followed by (b) a 12-day acquisition (Acq) phase, in which two sucrose pellets were administered 10 seconds after the retraction of the lever assigned as CS+, but not the lever on the other side (CS-), independent from the animals’ response. After acquisition, the next stages included (c) 12 days of reversal (Rev), swapping CS+ and CS– levers, (d) 6 days of extinction (Ext) to disassociate the rewards with the levers, (e) 6 reinstatement (RI) sessions with the reintroduction of rewards, and (f) 6 days of negative automaintenance (NA), where rats were only rewarded when they refrained from pressing the CS+ lever. Sign-tracking behaviors were quantified with a PCA index, calculated as an average of three metrics related to the differences in interactions with the levers and food magazine, as described previously,^75^ where higher PCA indices indicated greater sign-tracking tendencies. Additional measures of sign-tracking, including probability and latency to press the CS+ or CS– lever, or interact with the food magazine were recorded.

### Tail Flick Test

As previously,^72^ thermal pain sensitivity was assessed using a Columbus Instruments Tail-Flick Analgesia Meter. This device included a continuously illuminated shutter-controlled lamp (8V high intensity (6 amps)) as the heat source, which was set to an intensity level of 10 for this test. Homo (n = 20), Het (n = 20), and WT (n = 23) rats were first wrapped in a towel to prevent unwanted movement. Their tails were placed above the lamp. A timer was started once the lamp was turned on and was automatically stopped with the tail flick latency recorded once the rats lifted their tails to indicate a pain response.

### Statistical analyses (sign-tracking)

All statistical tests used α = 0.05. To assess differences in sign-tracking behaviors between sexes and genotypes, statistical analyses were performed using a session × sex (Male vs. Female) × genotype (Homo vs Het vs WT) mixed-design repeated measures ANOVA. Tukey post-hoc corrections were used following significant main effects of genotype or sex x genotype interaction effects. Data from the open field test and tail flick test were analyzed similarly for both sex and genotype effects using a sex x genotype two-way ANOVA.

### Pre-pulse inhibition and behavior pattern monitor

A total of 39 CADM2 littermates were identified as wildtypes (n=12; ♂=6 and ♀=6), heterozygotes (n=16; ♂=7 and ♀=9), or full knockouts (KO; n=11; ♂=9 and ♀=2) of for the CADM2 gene mutation. Rats were housed in dyads in clear plastic containers in a climate-controlled room on a 12-hour light/dark schedule (7:00 AM-7:00 PM dark). Operant training commenced at ∼12 weeks of age. Operant-trained rats were maintained at ∼90% of their free-feeding body weight. Water was available *ad libitum*, except during training and testing. The 32 additional non-operant trained rats were not food restricted at any time, while operant-trained rats were not food restricted when tested in PPI or BPM. Training and testing occurred during the dark portion of rats’ light/dark schedules. Rats were maintained in a dedicated animal facility compliant with all federal and state requirements and approved by the American Association for Accreditation of Laboratory Animal Care.

Non-food deprived rats were placed into startle chambers to assess sensorimotor gating using the acoustic startle response and pre-pulse inhibition (PPI) paradigm. A week later, animals were tested in the Behavioral Pattern Monitor (BPM) to measure locomotor and exploratory behavior. Detailed methodology and statistical analyses for PPI and BPM tests have been previously described in full detail.^76^ After BPM testing, subjects were food-deprived to 85% baseline body weight and trained to make a nose poke response on a fixed-ratio 1 schedule of reinforcement (FR1) in preparation for testing in the Iowa Gambling Task (IGT; measures risk-based decision-making), Probabilistic Reversal Learning Task (measures reinforcement learning and reversal learning), and the Progressive Ratio Breakpoint Task (PRBT; measures motivation). Once trained, animals were tested in each task once, with FR1 training sessions interleaved between each test. Detailed methodology and statistical analyses for these tasks have been previously described in full detail.^77,78^ Where sphericity violations were found, Greenhouse Geisser corrections were appropriately applied.

### Online resources

Scripts are deposited at https://github.com/talkowski-lab/cadm2_recursive_splicing. RNA-seq data will be deposited at dbGaP.

## Notes

### Competing Interest Statement

The authors have declared no competing interest.

