## Supplemental Figures for "Aberrant recursive splicing in a human disease locus"

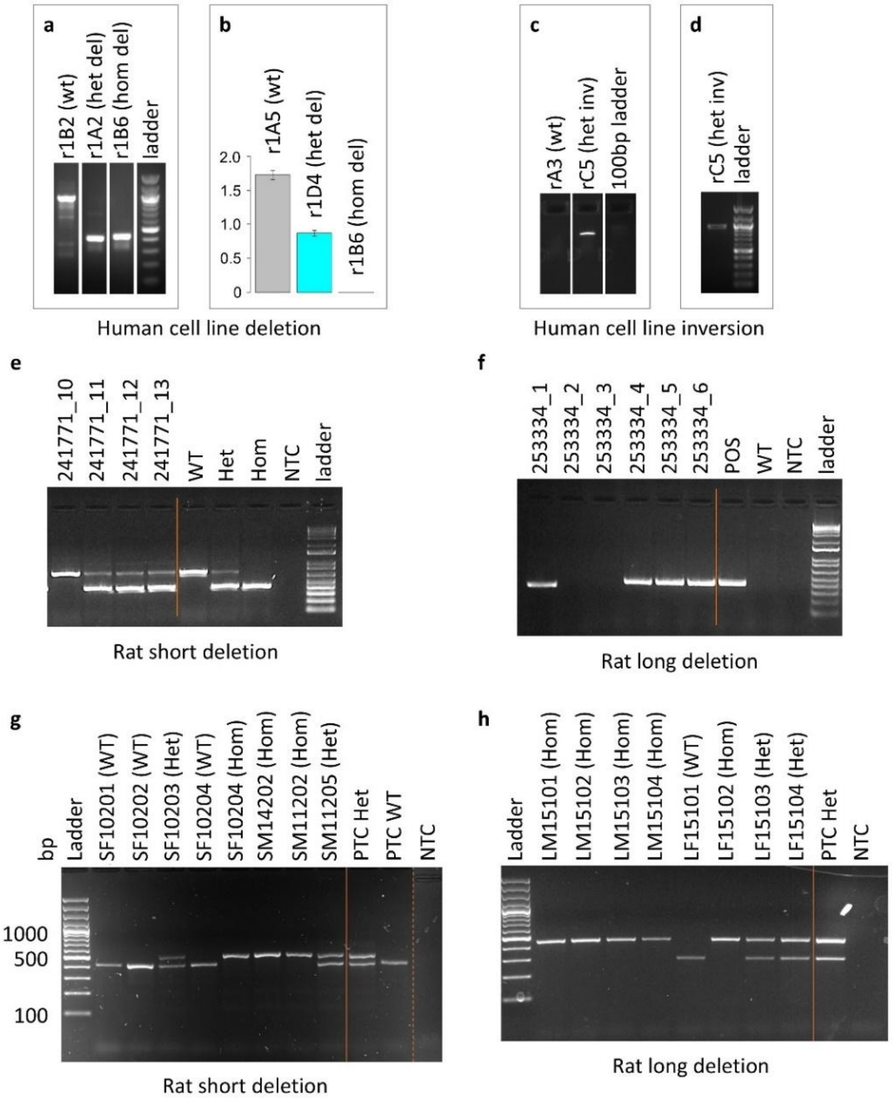


**Figure S1. Example genotyping of *CADM2* RS1 deletions and inversion in CRISPR engineered iPSCs and gene-edited rats. a.** PCR to detect the ~500 bp RS1 deletion. Del, deletion. Het, heterozygous. Hom, homozygous. **b.** ddPCR to detect and determine the zygosity of RS1 deletions, both 500 bp and 100 kb. Scale is approximate ploidy. **c.** PCR to detect the 500 bp RS1 inversion. Inv, inversion. **d.** PCR spanning one breakpoint of the 500 bp RS1 del/inv region, which yields a product if one wt copy is present. This demonstrates that the inversion in rC5 is heterozygous. **e-f.** Rat RS1 del genotyping (approach 1). For each type of deletion (short or long), a single PCR with deletion-spanning primer set was run. **e.** PCR to screen for the 322 bp (short) deletion in edited rats. Lane 1 has no deletion. Lanes 2-4 are heterozygous. Lanes 5-7 are positive controls (wt, het del, and hom del, respectively). NTC, no template control. **f.** PCR to screen for the 85 kb (long) deletion in edited rats. Lanes 1, 4, 5, and 6 have a deletion. Lanes 2 and 3 do not have a deletion. Lanes 7-9 are positive and negative controls (deletion (POS) and wt, respectively). NTC, no template control. **g-h.** Rat RS1 del genotyping (approach 2). For each type of deletion (short or long), one PCR with deletion spanning primers and another PCR that amplifies only the undeleted allele were pooled for each well. **g.** PCR to screen for the 322 bp (short) deletion in edited rats. PTC are positive controls (WT and Het). NTC, no template control. The reactions were performed separately, and their products were subsequently combined and analyzed by gel electrophoresis. **h.** PCR to screen for the 85 kb (long) deletion in edited rats. PTC are positive controls (WT and Het). NTC, no template control.

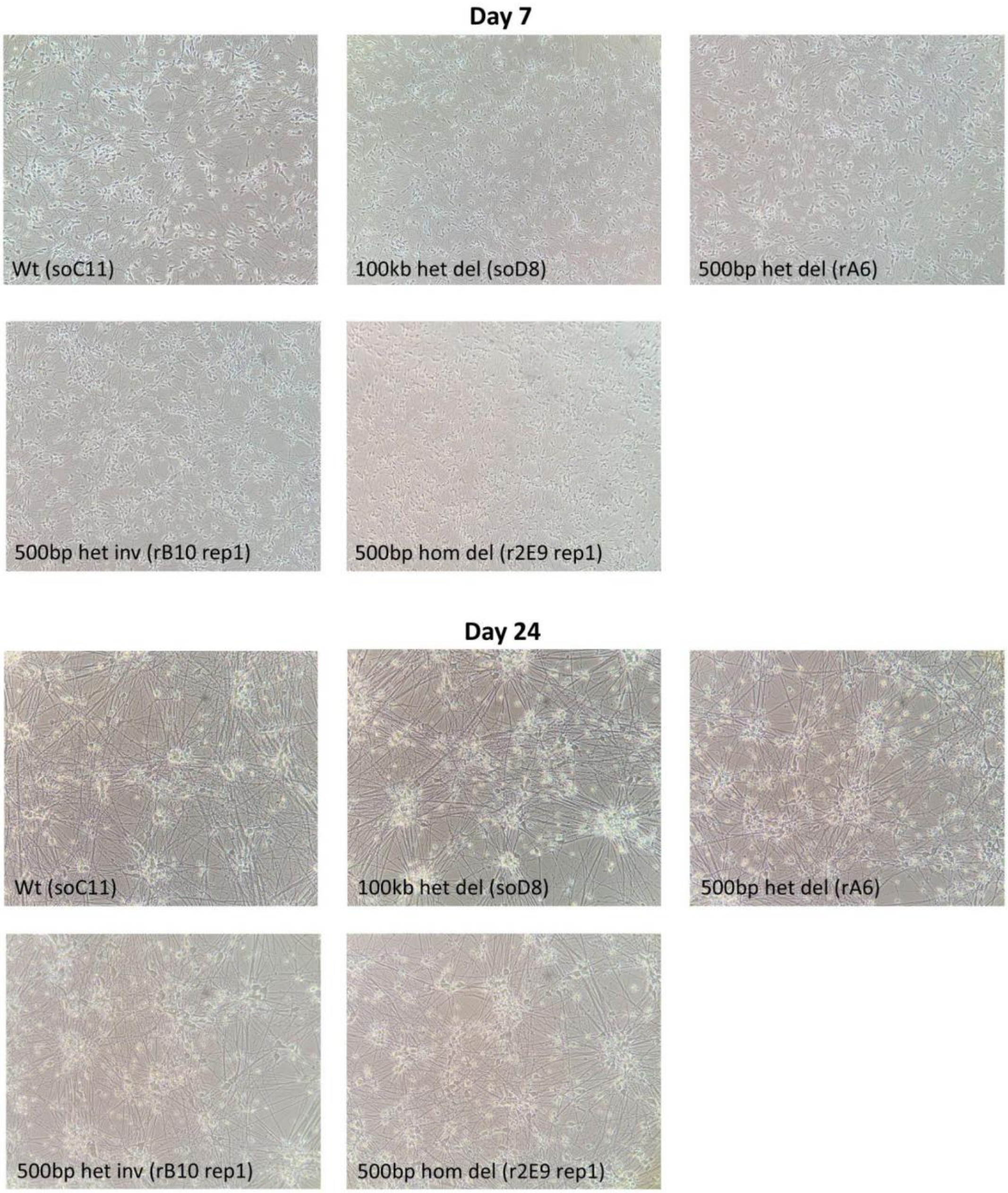


**Figure S2. Example induced neuron morphology.** One example cell line is shown for each genotype, early in differentiation (day 7) and at the end of differentiation (day 24).

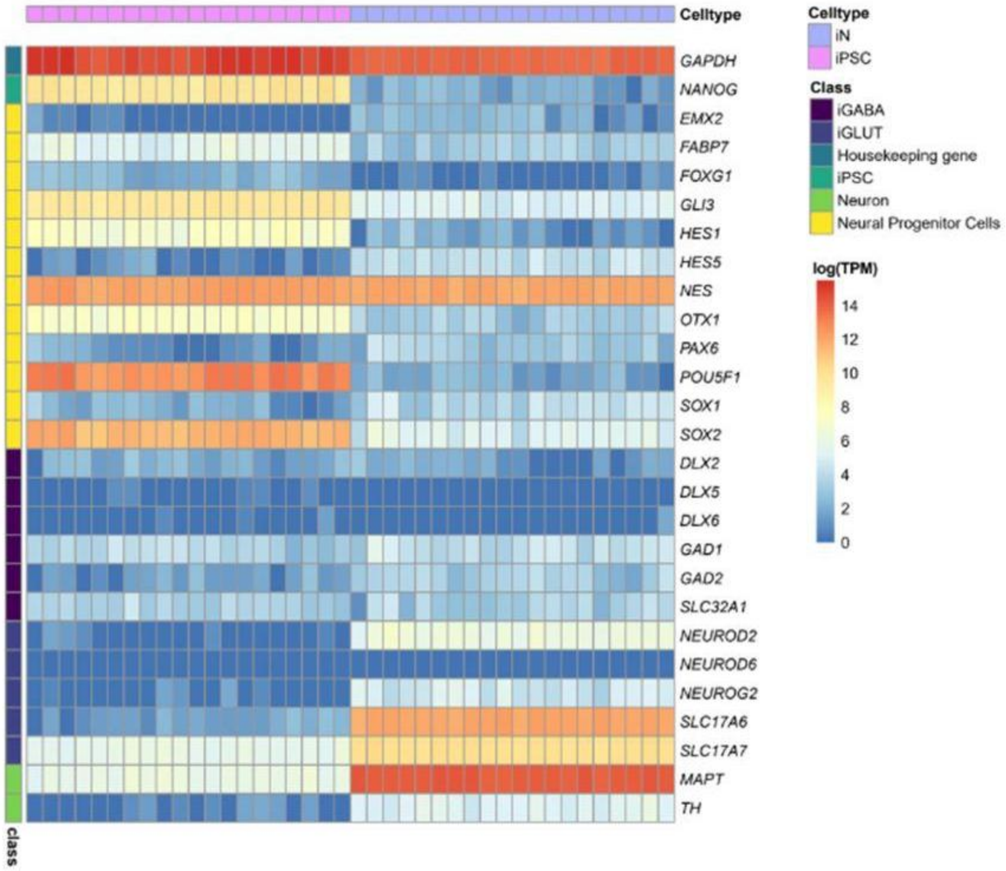


**Figure S3. RNA-seq quality control.** Relative expression levels of selected genes within several classes (iPSC, GABAergic, glutamatergic, neural progenitor, neuron, and housekeeping control) confirm that iPSC models reflect the pluripotent state and iN models reflect the neuronal state.

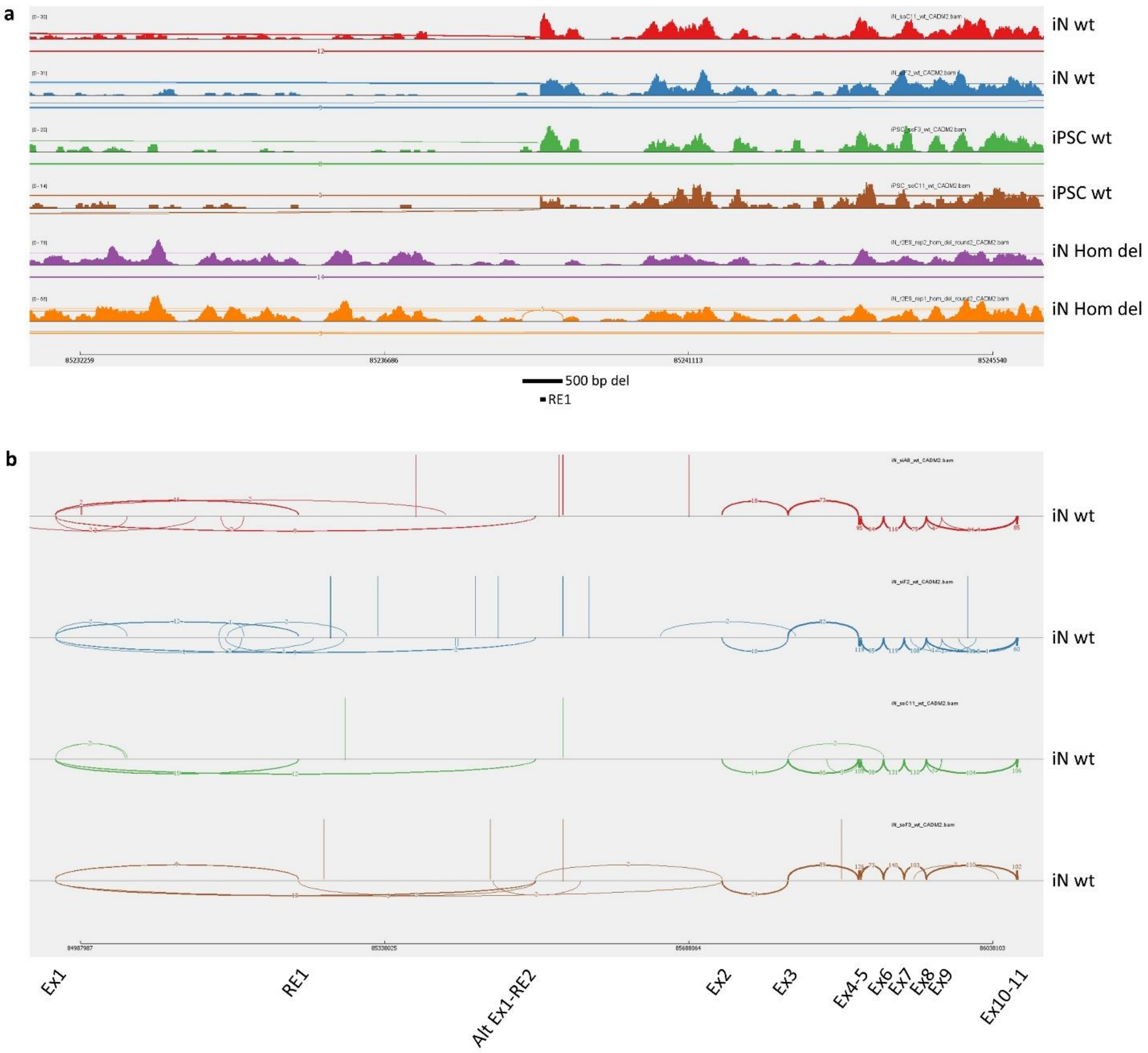


**Figure S4. Sashimi plots. a.** Localized read plots showing coverage of the recursive exon 1 (RE1) in wt iNs and iPSCs, and the loss of this coverage in hom 500 bp del iNs. The abundance of intronic sequence preceding RE1 is increased in hom del samples, likely by a failure to remove by splicing (see Figs. 2d, S9d). **b.** Sashimi plot of the entire *CADM2* locus, in wt iNs, with exons and predicted RS sites labeled. Splicing from exon 1 to RE1 and to RE2 are noted.

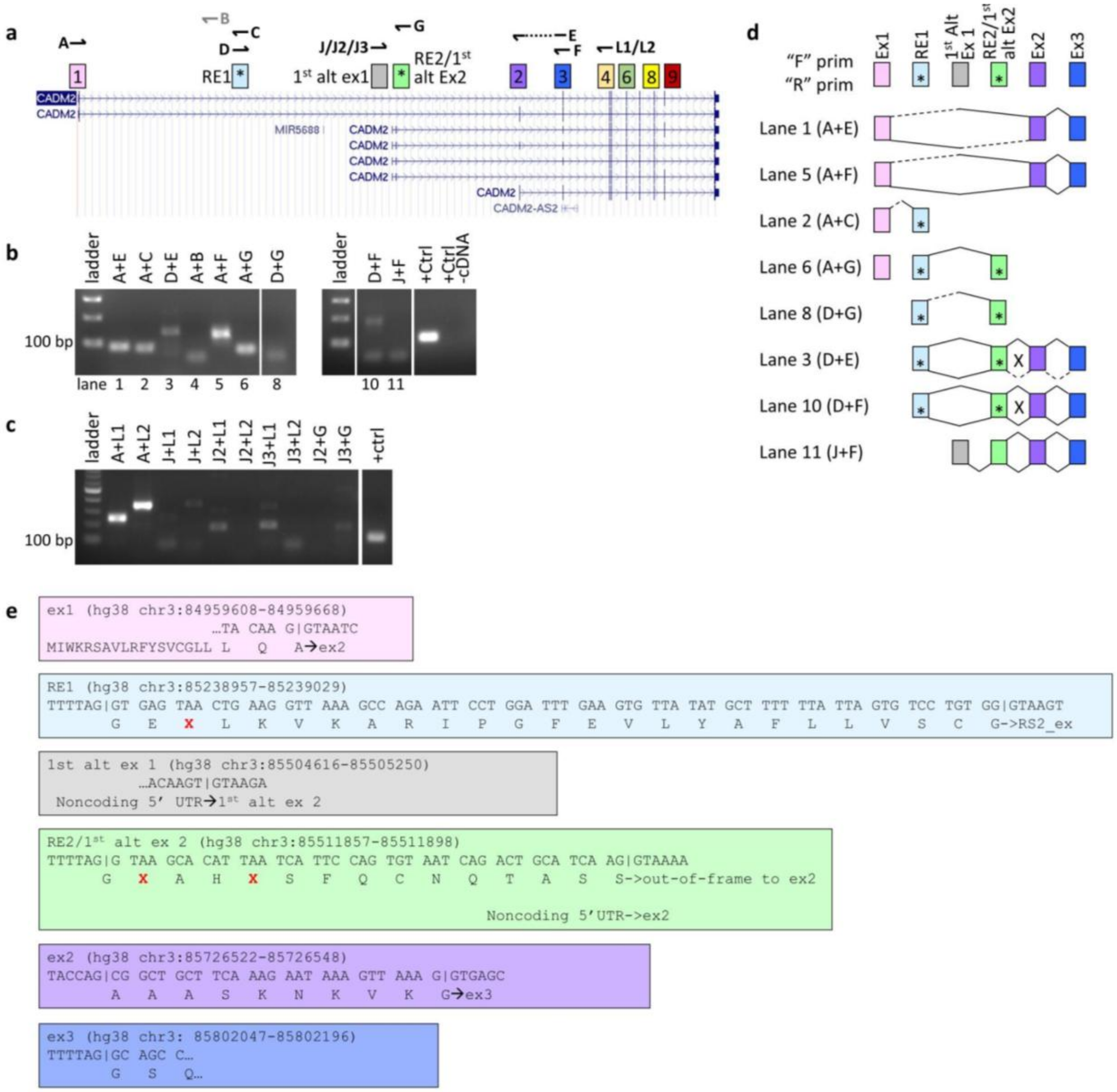


**Figure S5.** ***CADM2*** **isoform census in human brain RNA. a.** RT-PCR primers (half-arrows) above key exons (boxes with exon numbers) and the UCSC Genes track of the UCSC Genome Browser. A dotted line indicates that a primer spans two exons. **b.** RT-PCR of cDNA, to generate amplicons for Sanger sequencing (d). Lane numbering is from source gel and to ease interpretation of panel (d). A lack of band in lane 4 (primers A+B) is expected, as B is proximal to recursive exon 1 (RE1). **c.** RT-PCR of cDNA using additional primers confirms that inclusion of the 1st alternative exon 1 (1st alt ex 1) is rare, as no robust bands are produced by any of three forward primers in that exon in combination with reverse primers in the 1st alt ex 2 or in exon 4. **d.** Splice junctions detected by Sanger sequencing of RT-PCR products in (b) (solid splice lines) or inferred by RT-PCR but unable to be completely resolved by sequencing owing to proximity to the primer (dotted splice lines). Several unannotated transcripts are identified. * = Stop codons. X = out-of-frame junction. Splice lines denoting splicing that connect the tops of exons are derived from forward ("F") primer-derived sequences, and those below the line are from reverse ("R") primer-derived sequences. Half-solid/half-dotted splice lines indicate that the whole exon on the solid side was sequenced. **e.** Predicted translational outcomes of Sanger-validated transcripts, through exon 3. Coloring of boxes matches exon coloring in (d). DNA sequences are shown above predicted amino acid sequences. | = splice junctions. Red X's are stop codons. The coordinates for exon 1 are the coding portion of the exon. Coordinates for the start of exon 1, the start of alt 1st ex 1, and the end of exon 3 are via the UCSC Genes track of the UCSC Genome Browser.

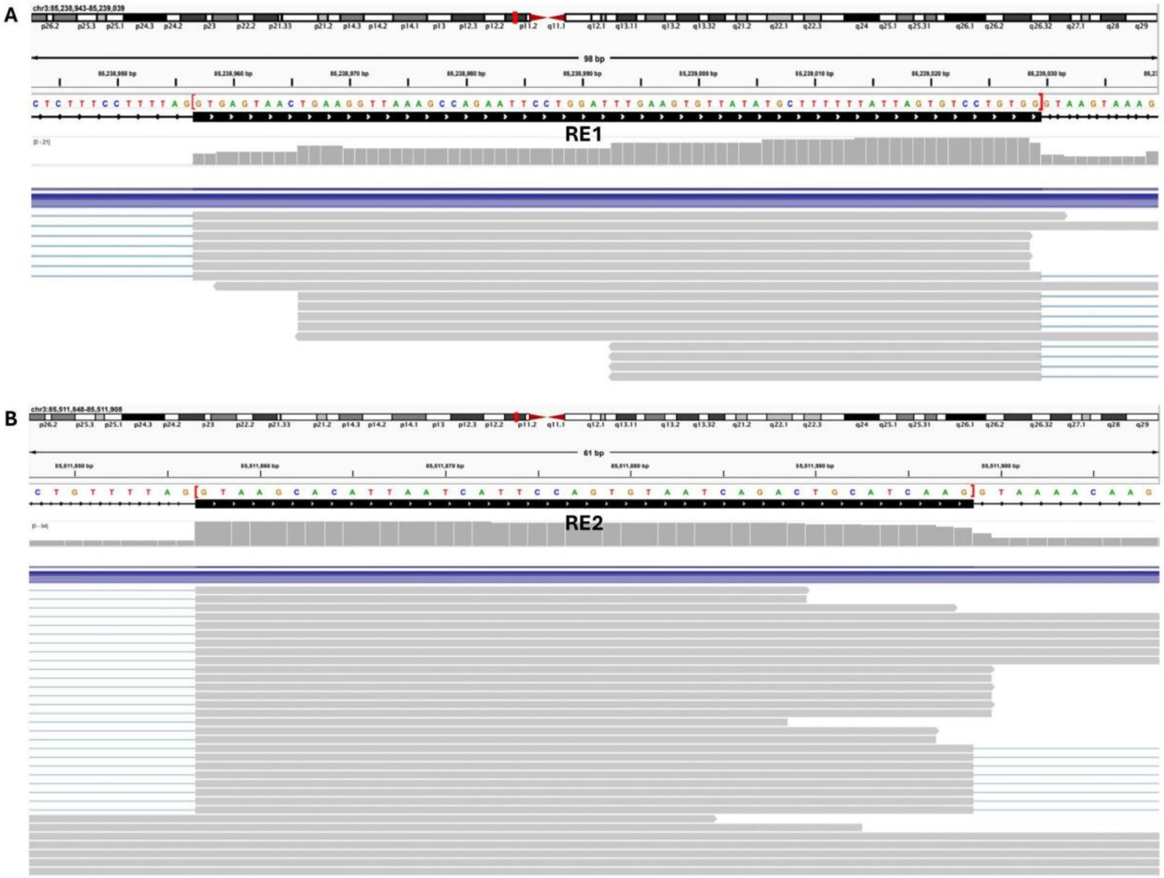


**Figure S6.** ***CADM2*** **RE1 and RE2, as defined by total RNA-seq in human iNs. a.** RE1. Red brackets show the extent of the RE, which matches (to the base pair) with RE1 as defined by RT-PCR (Fig. S5). **b.** RE2. Red brackets show the extent of the RE, which matches (to the base pair) with RE2 as defined by RT-PCR (Fig. S5).

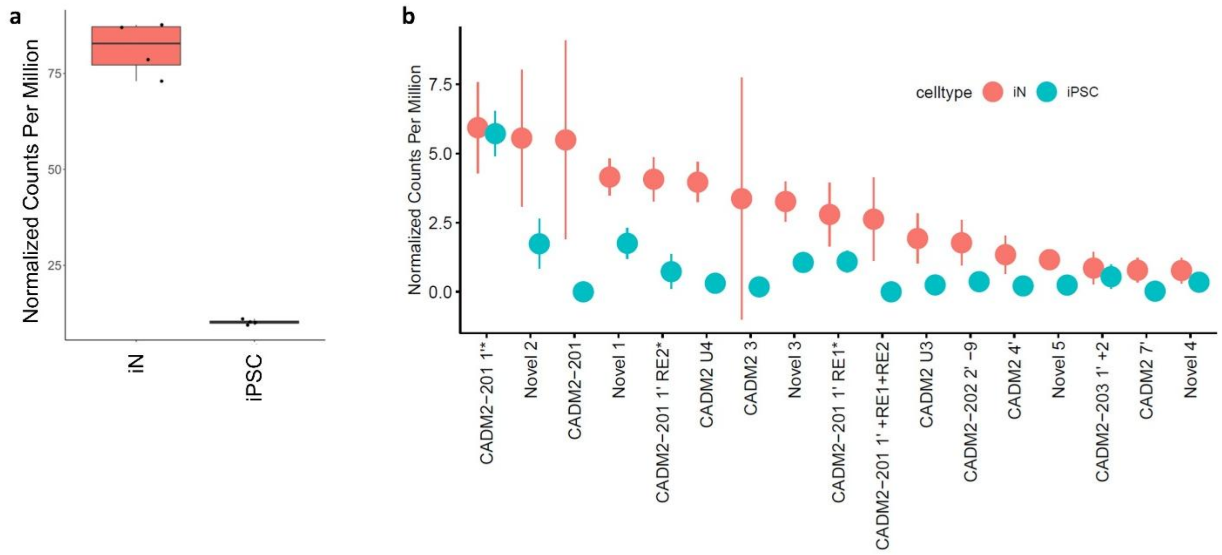


**Figure S7. Differences in *CADM2* expression between iPSCs and iNs. a.** Corrected gene expression showing that *CADM2* is expressed at a substantially lower level in iPSCs. **b.** There is lower overall expression and a dramatic difference in *CADM2* transcript usage between iPSCs and iNs (see Fig. 3b). Data are normalized CPM.

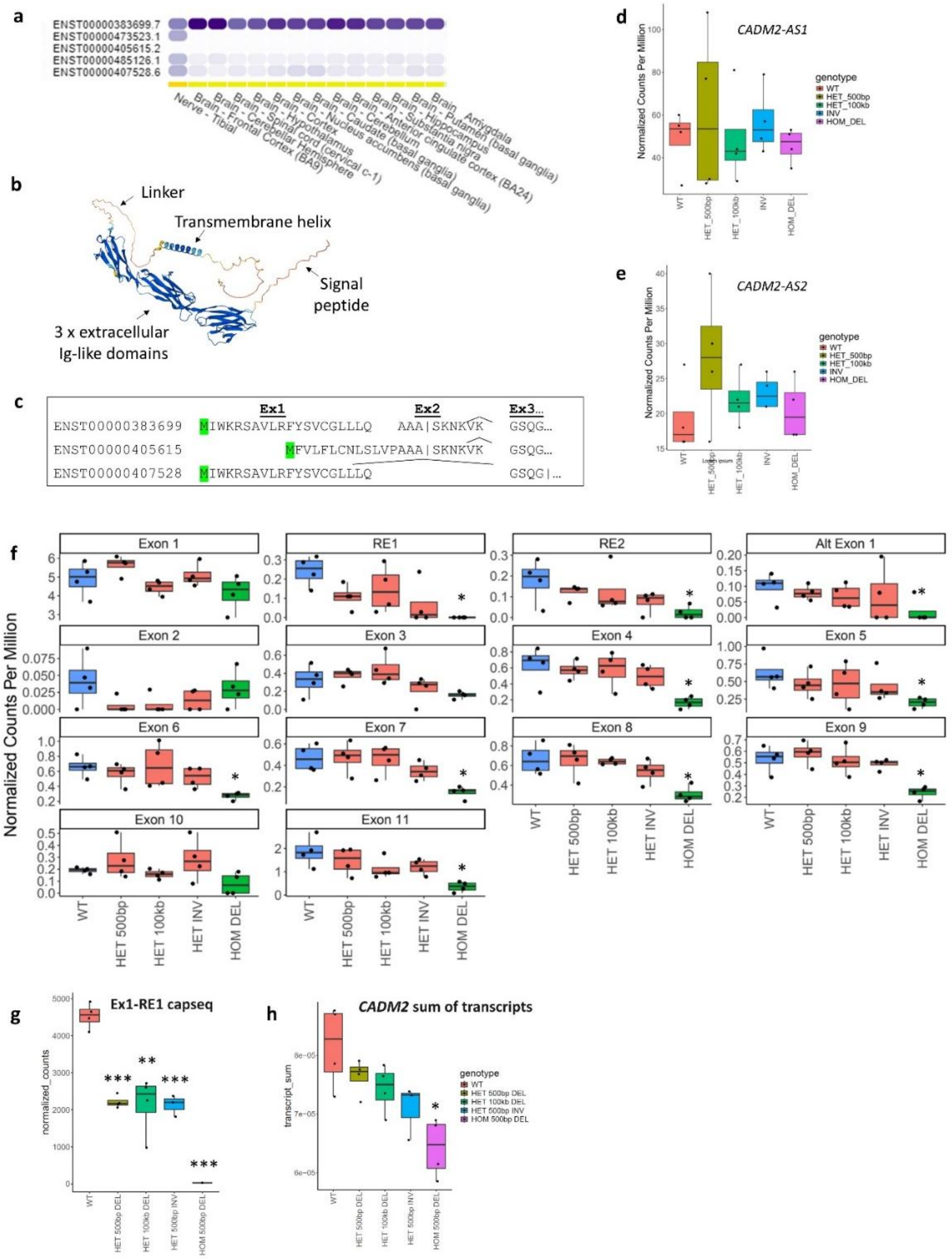


**Figure S8. Exon, junction, and transcript expression, extended. a.** *CADM2* isoform expression data from GTEx, showing only neuronal tissues. **b.** CADM2 AlphaFold Protein structure (via UniProt https://www.uniprot.org/uniprotkb/Q8N3J6/entry). Inclusion of exon 9 is expected to increase the length of the linker between CADM2's transmembrane helix and extracellular Ig-like domains. **c.** Differences among transcripts' signal peptide completeness. | indicates the position of predicted signal peptide cleavage. Signal peptide scores (via Signal P 5.0; https://services.healthtech.dtu.dk/service.php?SignalP-5.0) are also predicted to differ among isoforms (not shown). Exon 2 (in-frame) encodes part of the signal peptide. Exon 9 (in-frame) encodes much of the linker between the extracellular Ig domains and the transmembrane helix. **d.** Expression of *CADM2-AS1* in iNs, by genotype, showed no significant differences. **e.** Expression of *CADM2-AS2* in iNs, by genotype, showed no significant differences. **f.** iPSC exon expression, as boxplots by genotype. See Table S12 for statistics; displayed here FDR. * FDR<0.1. **g.** Exon1-RE1 junction from Cap-seq data, showing that recursive splicing is ablated in a zygosity-dependent manner upon deletion of RE1. **h.** Summing the *CADM2* transcripts in Fig. 3b by genotype identifies a 21.2% decrease in homozygous iNs (p=8.8e-3). Significance markings are * p<0.05, ** p<0.005, *** p<0.0005.

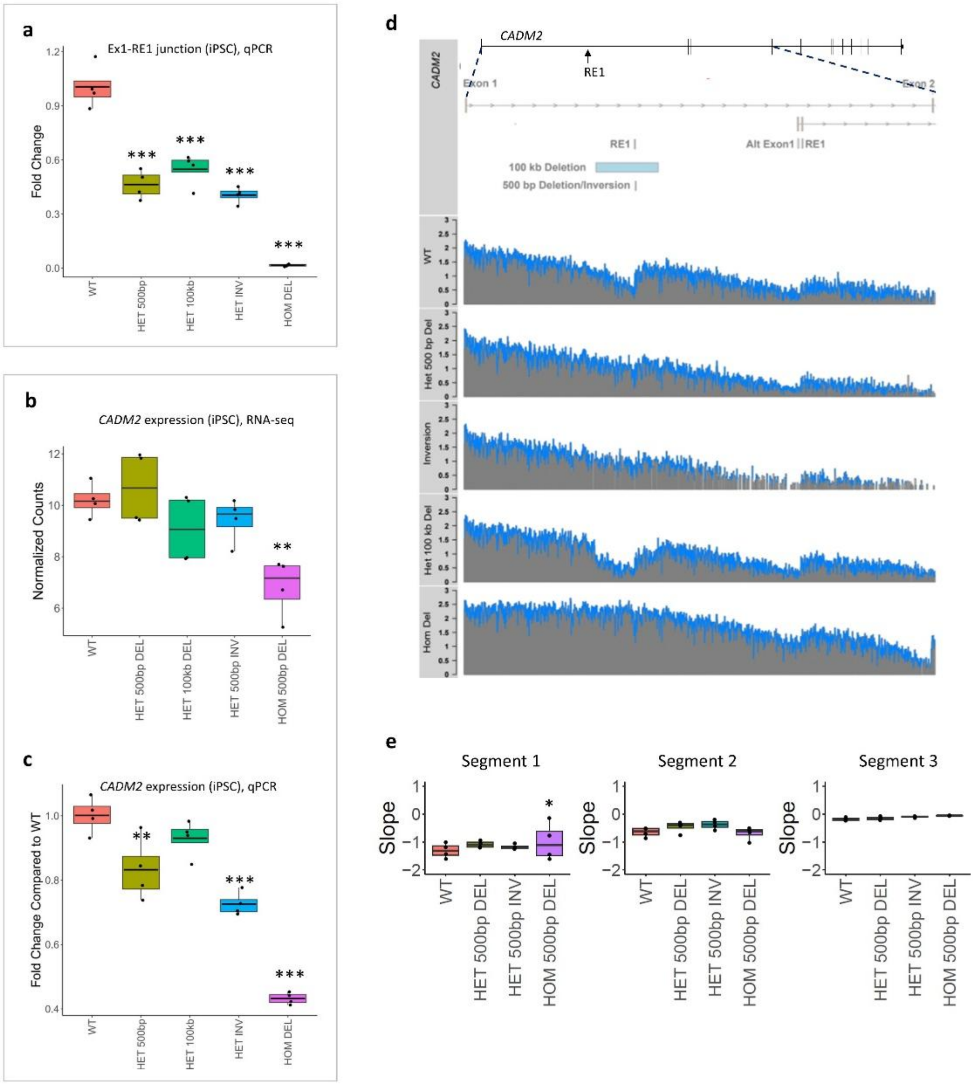


**Figure S9. CRISPR *CADM2* RS1 site deletion in human induced pluripotent stem cells (iPSCs) ablates recursive splicing, alters the pattern of intronic mRNA abundance, and decreases *CADM2* expression. a.** qPCR using primers amplifying from exon 1 to RE1 confirms that RS1 deletion ablates recursive splicing in a zygosity-dependent manner, with a decrease of 45-59% in heterozygous and of 98% in homozygous lines, compared to wt (one-way ANOVA p=6.25e-10; all comparisons to wt p<1e-5 by Tukey's HSD test). Normalization gene is *GUSB*. **b.** RNA-seq demonstrates that RS1 site deletion decreases the overall expression of *CADM2*. The decrease is 37.9% in homozygous iPSCs compared to wt (p=5.9e-5 by Wald's test). **c.** qPCR using primers amplifying between constitutive exons 6 and 8 confirms that RS1 site deletion decreases expression of *CADM2* in a dose-dependent manner, by 7-27% of wt in heterozygous and by 57% in homozygous lines (one-way ANOVA p=7.59e-9; individual comparisons to wt significant for hom 500 bp del (p<1e-6), het 500 bp del (p=0.0083), and het 500 bp inv (p=7.69e-5), by Tukey's post-hoc HSD test, as well as when hets are combined (p=1.1e-2)). Normalization gene is *GUSB*. **d.** Normalized total RNA-seq coverage ($log_2$ reads/bp) mapped across *CADM2* intron 1 shows the sawtooth pattern of read depth corresponding to known waypoints of recursive splicing (RS1, RS2). RS1 deletion or inversion alters the sawtooth pattern. Blue represents smoothened data. **e.** Read count slopes (normalized read depth/cumulative relative position), by genotype, within the segments defined in Fig. 2e. Ablation of RS1 flattens the slope in segment 1, significant for hom 500 bp del (p=3.27e-2 by Student's t-test). Het 100 kb del was removed from the analysis on account of removing a large part of the intron. Significance markings are * p<0.05, ** p<0.005, *** p<0.0005.

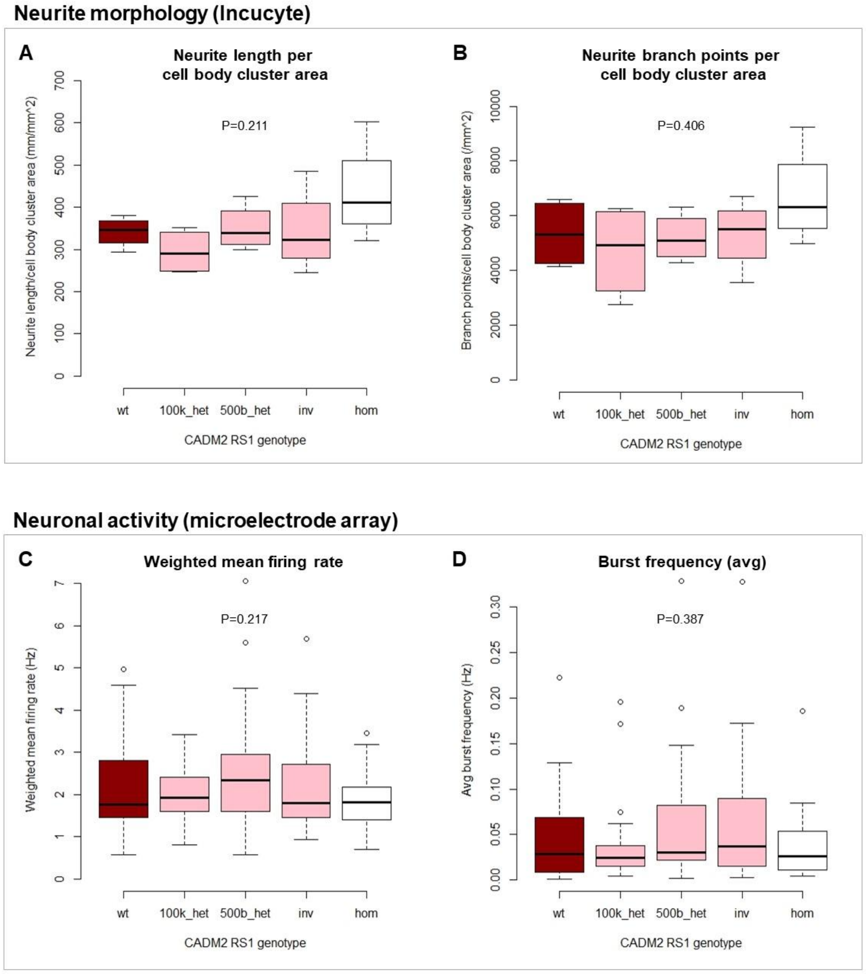


**Figure S10.** ***CADM2*** **RS1 deletion does not alter gross neurite morphology or neuronal activity. a-b.** IncuCyte live cell analysis of induced neurons (iNs) at time point 2 (days 18-20 of differentiation), 10k cells/well. Neither the neurite length per cell body cluster area nor the number of neurite branch points per cell body cluster area differed among genotypes. **c-d.** Microelectrode array analysis at day 27 of differentiation, filtered for wells with ≥8 active electrodes. Weighted (adjusted for the number of active electrodes) mean firing rate. Average (avg) burst frequency is the total number of single-electrode bursts/duration of analysis. Wt lines are in red. Het 100 kb del (100k_het), het 500 bp del (500b_het), and het 500 bp inv (inv) lines are in pink. Hom 500 bp del (hom) lines are in white. Statistics are one-way ANOVA.

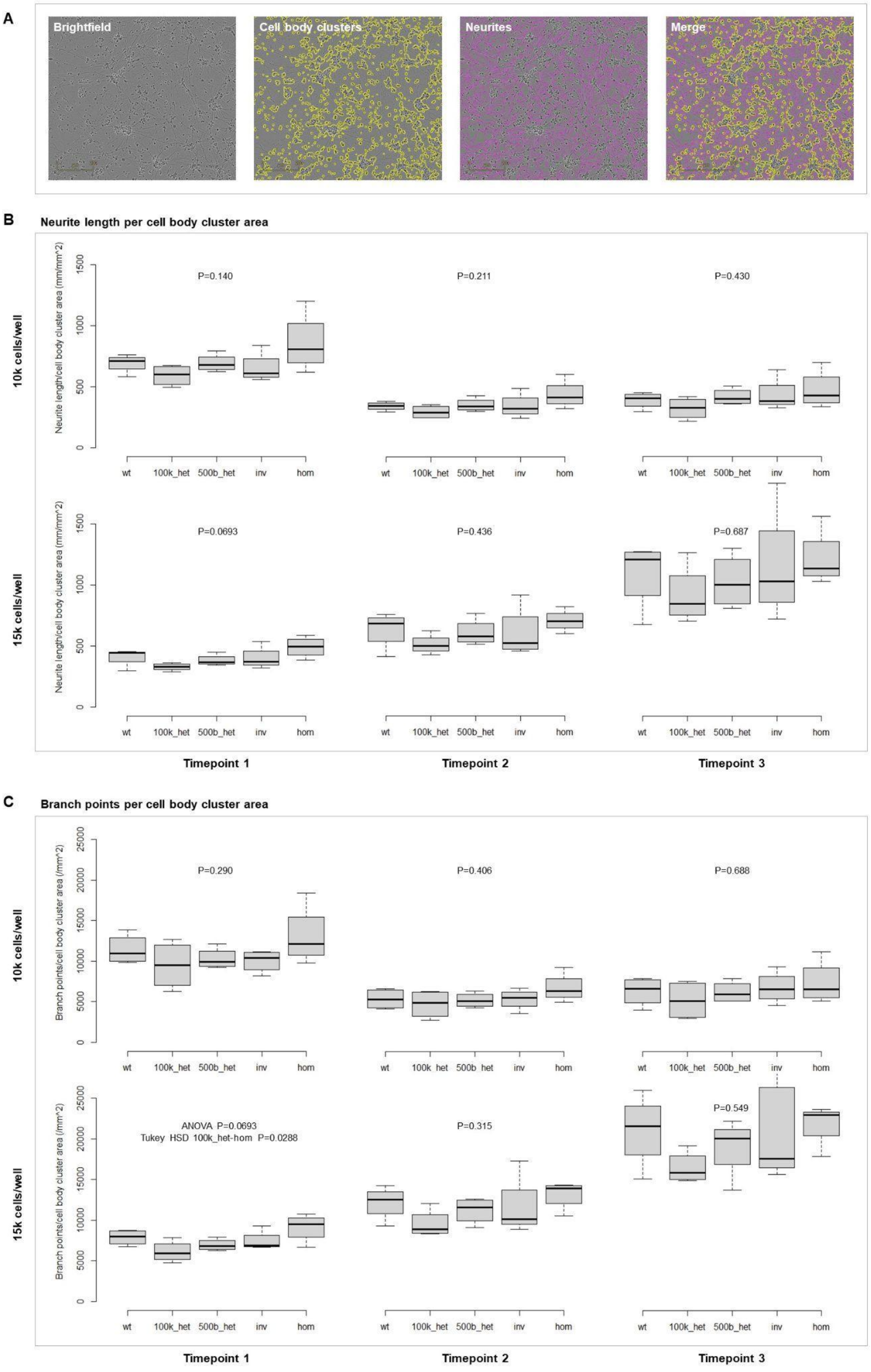


**Figure S11. Extended IncuCyte live cell analysis data.** *CADM2* RS1 deletion does not alter gross neurite morphology via any of an extended set of metrics. Two iN plating densities (10k and 15k cells/well) and three timepoints (1 = iN differentiation days 11-13; 2 = days 18-20; 3 = days 25-27) were assessed. **a.** Example of automated calling of cell body clusters and neurites, 10k cells/well, timepoint 3. **b.** Neurite length per cell body cluster area. **c.** Branch points per cell body cluster area. 100k_het, het 100 kb del. 500b_het, het 500 bp del. inv, het 500 bp inv. hom, hom 500 bp del. Statistics are one-way ANOVA.

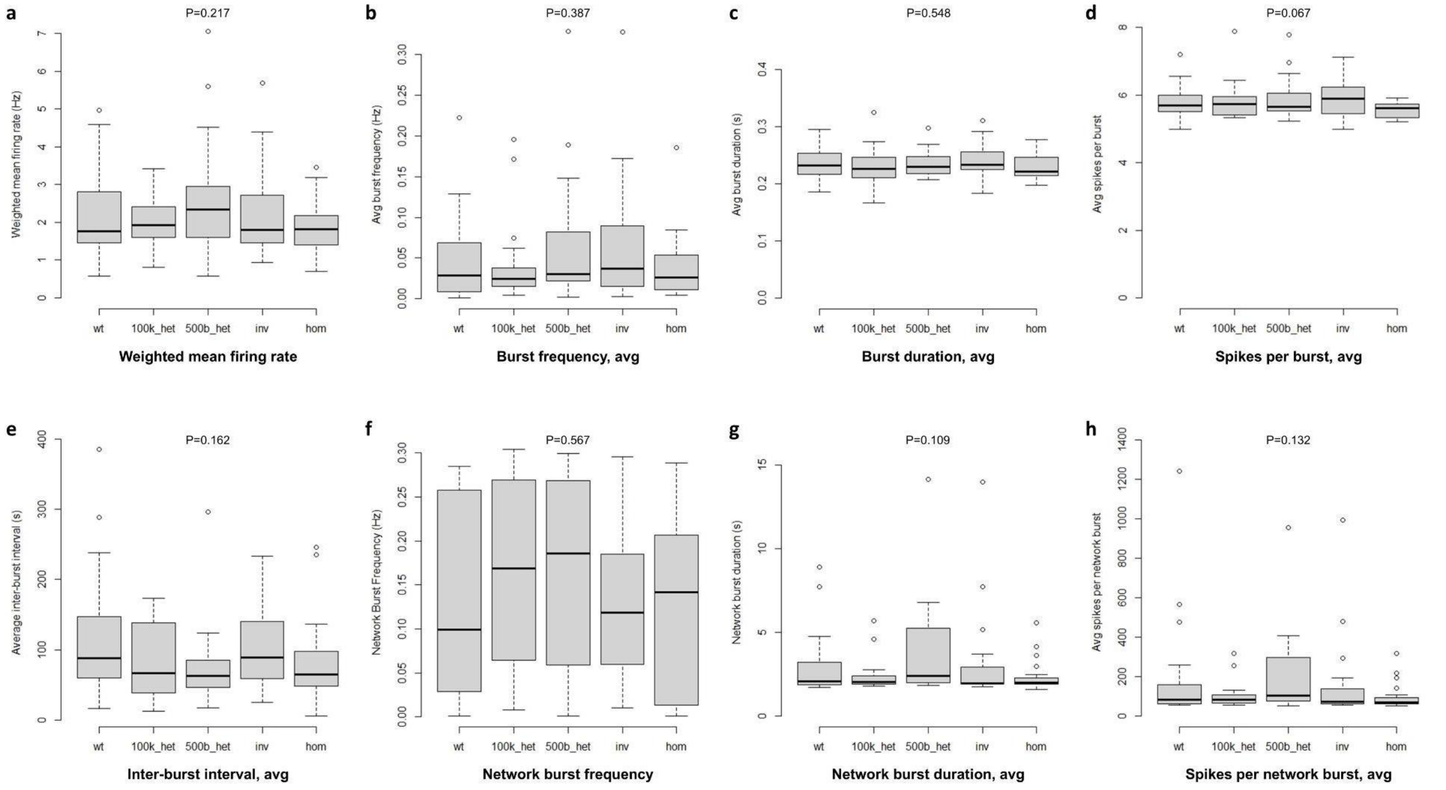


**Figure S12. Extended microelectrode array data.** Microelectrode array analysis at day 27 of differentiation, filtered for wells with ≥8 active electrodes. *CADM2* RS1 deletion does not alter neuronal activity via any of an extended set of metrics. **a-b.** Weighted mean firing rate and average (avg) burst frequency, reproduced from Fig. S10. **c.** Average burst duration. **d.** Average spikes per burst. **e.** Average inter-burst interval. **f.** Network burst frequency. **g.** Average network burst duration. Cropping of the y axis, for clarity, removed one outlier. **h.** Average spikes per network burst. 100k_het, het 100 kb del. 500b_het, het 500 bp del. inv, het 500 bp inv. hom, hom 500 bp del. Statistics are one-way ANOVA.

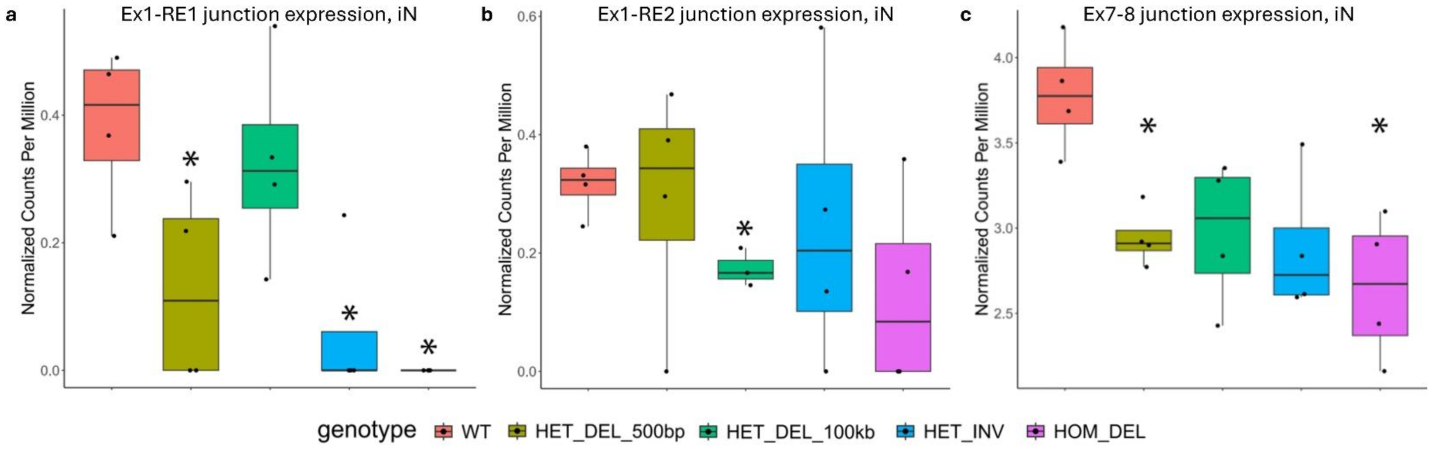


**Figure S13. Additional junction expression analyses, iNs. a.** Total RNA-seq splice junction analysis of the exon 1 (ex1) to recursive exon 1 (RE1) junction. The average reduction is by 55% across heterozygous iNs (down 66% in het 500 bp del, p= 0.04; down 14% in het 100 kb del, p=0.61; down 84% in het 500 bp inv, p=0.01; one-sided t-test) and by 100% in homozygous iNs (p=9.0e-3), compared to wt. The exon 1 to RE1 junction is rare (range of 6-12 junction spanning reads in wt iNs, as compared with 83-105 reads for the junction between constitutive coding exons 7 and 8). **b.** iN total RNA-seq junction from exon 1 (Ex1) to RE2 (het 100 kb del p=8.3e-3). **c.** iN total RNA-seq junction from exon 7 (Ex7) to exon 8 (Ex8) (het 500 bp del p=8.3e-3; hom 500 bp del p=6.7e-3).

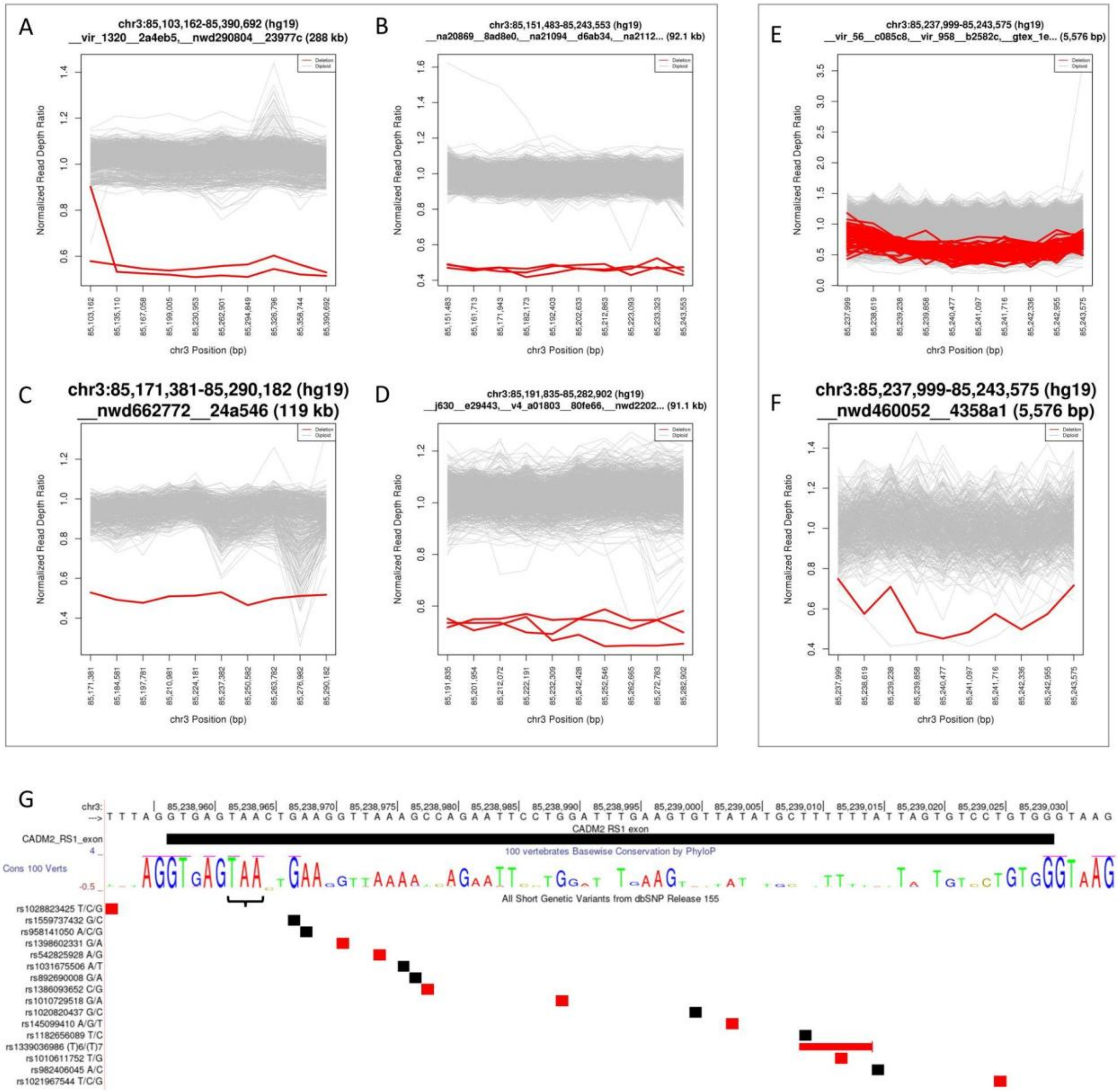


**Figure S14.** ***CADM2*** **RS1 deletions and simple nucleotide variants (SNVs) in gnomAD. a-d.** Read-depth plots showing the four intronic deletion alleles spanning *CADM2* RS1 in gnomAD v4. Red indicates non-reference samples. **e-f.** A smaller, ostensibly common 5.6 kb gnomAD deletion allele encompassing *CADM2* RS1 is a likely technical artifact; while individual batch analyses suggest outliers (f), the overall read depth distribution of non-reference samples is not distinctly different from the distribution of reference read depth (e) and most carriers are in either PCR+ batches or batches of high dosage bias (not shown). **g.** SNVs localized to the *CADM2* recursive exon 1 (RE1). Those in red are in gnomAD, while those in black are other annotated dbSNP (v155) variants. See also Table S9. All gnomAD SNVs are rare and no gnomAD or dbSNP SNVs affect the conserved splice sites or stop codon (bracket) of RE1 (black bar).

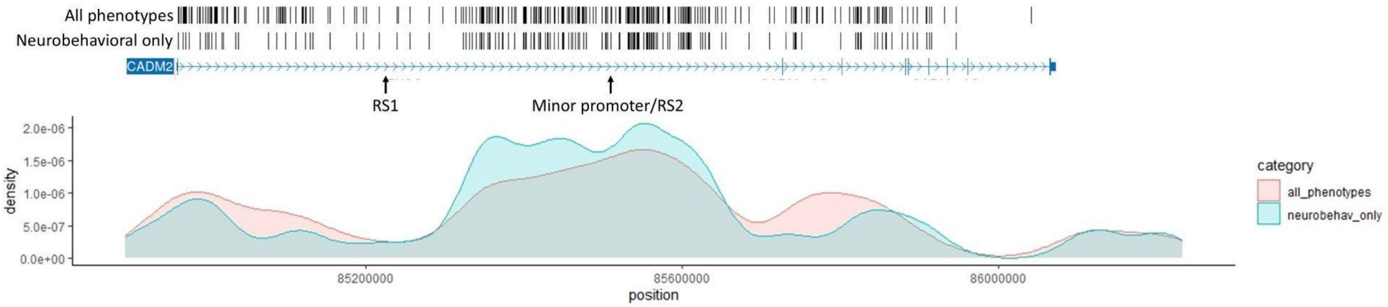


**Figure S15. GWAS hits in *CADM2* are overrepresented in the region of a minor promoter and the second recursive splice site (RS2).** Position of *CADM2* GWAS hits (https://www.ebi.ac.uk/gwas/) are shown as tick marks, for all phenotypes and restricted to neurobehavioral phenotypes (psychiatric, behavioral, substances, and cognitive/educational attainment). A density plot of each is shown below. Position is hg38 chr3.

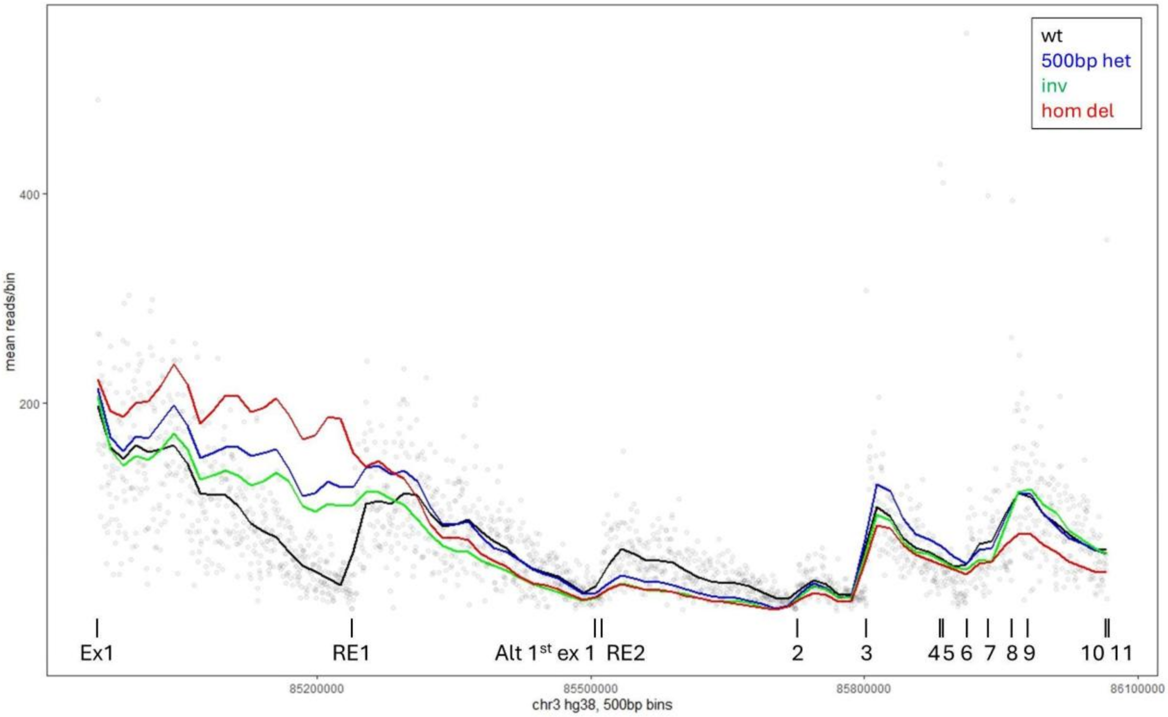


**Figure S16. Smoothened read depth over *CADM2* in iNs, by genotype.** The flattened slope of the mutant genotypes over the exon 1 (Ex1) to RE1 interval is apparent. The mildly higher slope in the RE1-RE2 interval in 500 bp het (Fig. 2f, "Middle Segment") is apparently attributable to a mildly higher starting read depth in that intronic interval. Dots are mean reads in wt per 500 bp bin, averaged across the four replicate lines. The trend lines are loess-smoothened curves, colored by genotype. Reads are normalized for library size per sample.

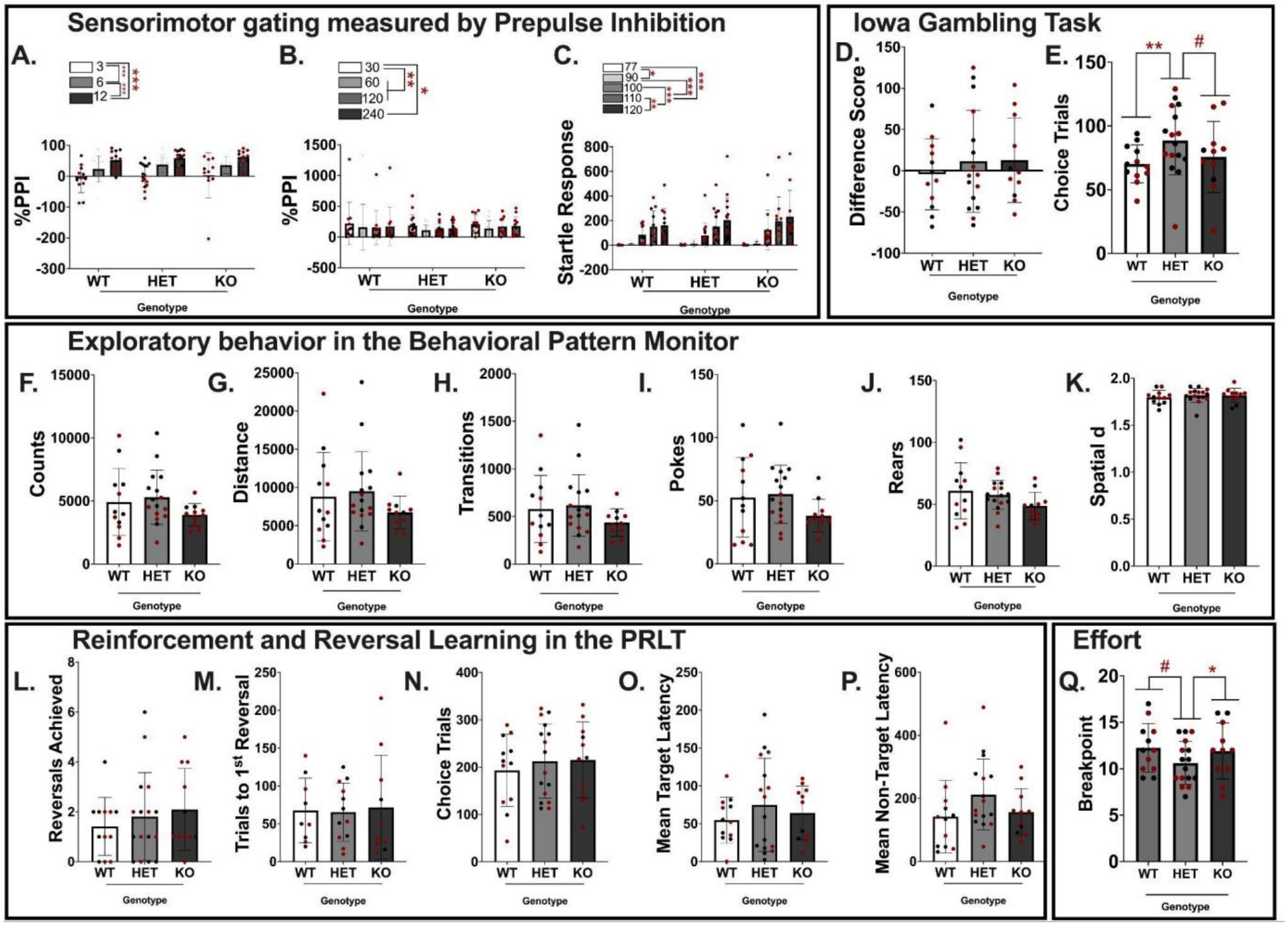


**Figure S17. Extended long del rat behavioral data. a-c.** *Cadm2* RS1 deletion did not disturb sensorimotor gating as assessed by pre-pulse inhibition (PPI) or startle response. **a.** Expected main effect of pre-pulse intensity on the percentage of PPI (%PPI) elicited [$F(1.343,44.313)=27.599, p<0.001$]; %PPI increased for all genotypes as the intensity of the pre-pulse increased (PP3 < PP6 < PP12, all $p$'s<0.001). **b.** Consistent with expectations, a main effect of interstimulus interval (ISI) length [$F(2.045,67.500)^*=6.347, p=0.003$] determined that the shortest ISI length produced higher %PPI (pre-pulse level=PP3) when compared to higher ISIs (ISI30 > ISI60, ISI120, ISI240, $p$'s<0.05-0.01). **c.** In addition, a main effect of pulse intensity [$F(1.616,53.326)^*=27.430, p<0.001$] determined that startle responses increased as the pulse intensity increased (P77 and P90 < P100, P110, and P120, $p$'s <0.001), indicating that rats of all genotypes similarly perceive and react to pulse tones. **d-e.** When risk-taking was measured using the Iowa Gambling Task, no impact of genotype was seen on difference score (d), but for the number of choice trials completed [$F(2,31)=4.445, p=0.020$] het completed more trials than wt rats ($p<0.01$) and tended to complete more trials than KO rats ($p=0.069$) (e). **f-k.** When exploratory behavior was assessed using the behavioral pattern monitor (BPM), no genotype effect on exploration was observed as measured by activity counts (f), total distance traveled (g), number of zone transitions (h), hole pokes (i), rearing (j), or spatial d value indicative of path linearity (k). **l-p.** Probabilistic reversal learning task (PRLT). No impact of genotype was seen on reversal learning (l) or reinforcement learning (m). The rats made the same number of choice trials (n) and showed no change in response latencies (o-p). **q.** When effortful motivation was assessed in the progressive ratio breakpoint test (PRBT) [$F(2,33)=3.480, p=0.043$], het rats tended to have lower breakpoints compared to wt rats ($p=0.096$) and had lower breakpoints than KO rats ($p<0.05$). Data presented as individual data points, mean, +/- S.E.M., with red and black individual data points for male and female data, respectively. ***$p<0.005$, **$p<0.01$, *$p<0.05$, #$p<0.1$ as indicated.

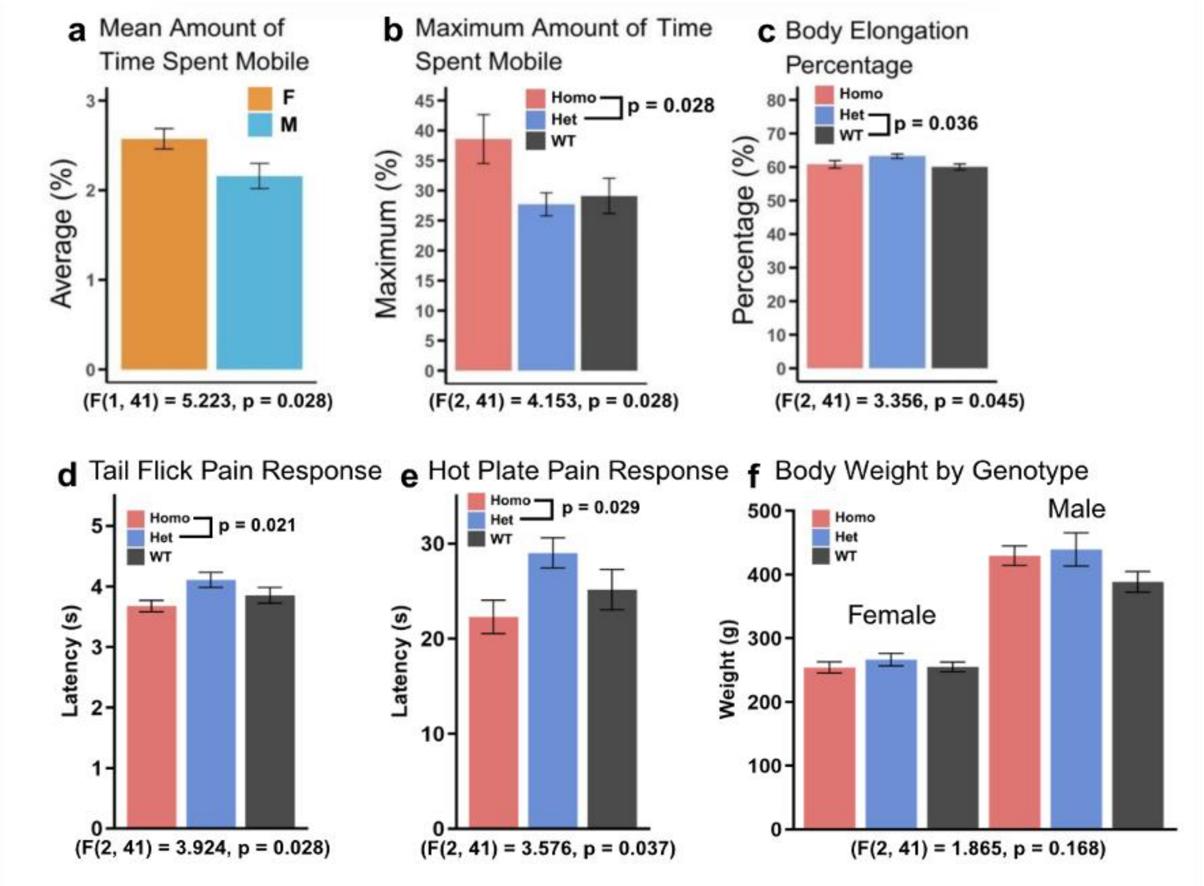


**Figure S18. Behavior and weight in *Cadm2* short del rats. a.** On average, female rats spent a significantly greater amount of time mobile compared to male rats in the Open Field Test. **b.** Hom short del rats spent a greater amount of time mobile compared to het short del rats. **c.** Het animals exhibited increased body elongation percentages compared to wildtype animals. **d-e.** Het short del rats displayed a significantly higher pain tolerance compared to hom short del rats in both the Tail Flick and Hot Plate behavioral paradigms. **f.** Body weights of animals did not differ by short del genotype (n=47; Wt: 8M, 8F; Het: 8M, 8F; Hom: 10M, 5F).

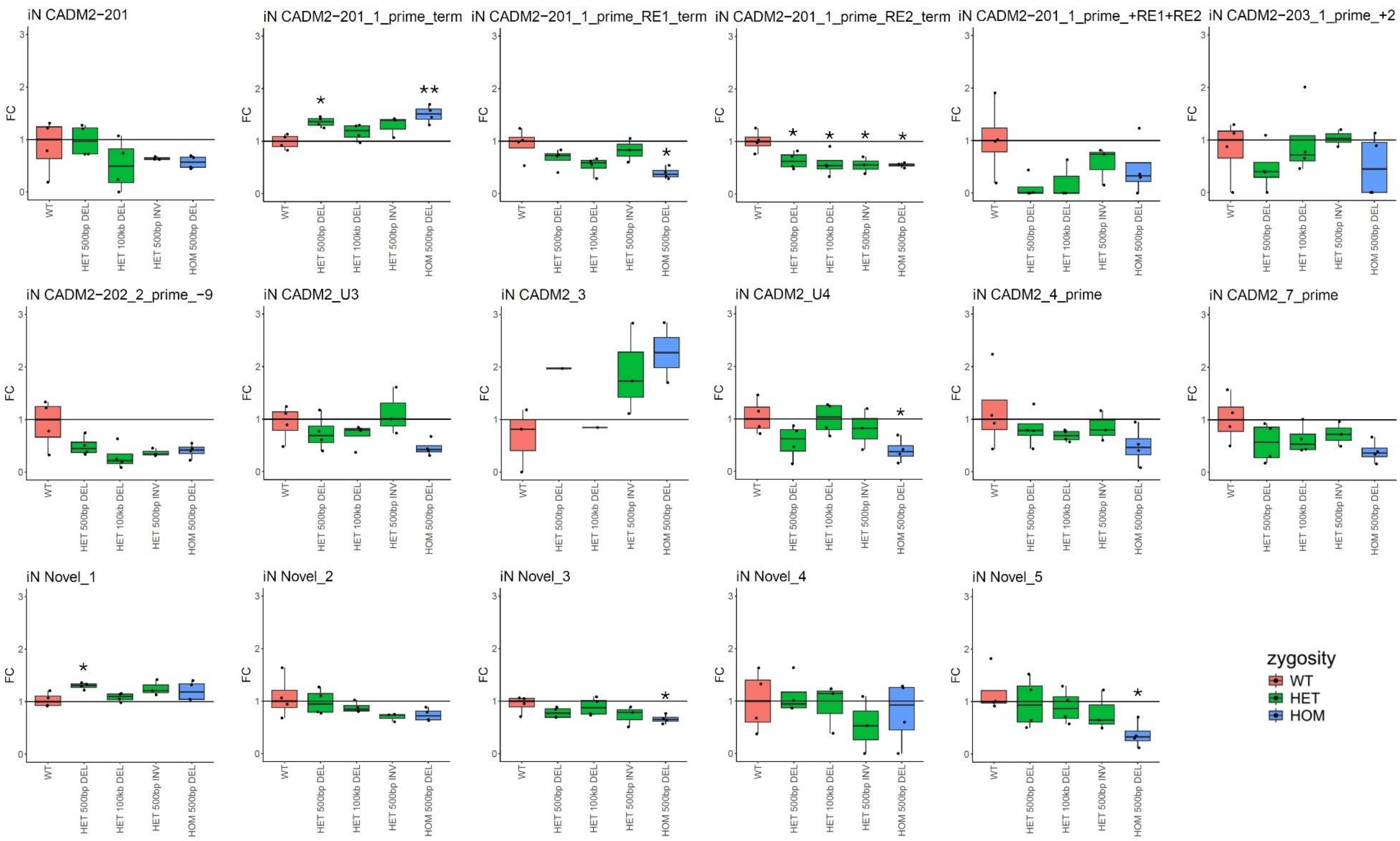


**Figure S19. Fold change of *de novo* reconstructed transcripts in iNs, extended.** Standard boxplots. Statistics are two-sided t-tests between the wt and edited cell line fold changes. * p<0.05, ** p<0.005, *** p<0.0005.
